# CXCL4-induced PBMCs Modulate Vascular Extracellular Matrix through Wnt5a-CaMKII-dependent Release of Calcific Extracellular Vesicles and Matrix Metalloproteinase-7

**DOI:** 10.1101/2023.05.15.540832

**Authors:** Jona B. Krohn, Florian Sicklinger, Anja Spieler, Susanne Dihlmann, Christian A. Gleissner, Hugo A. Katus, Norbert Frey, Florian Leuschner

## Abstract

**Background:** Macrophage heterogeneity plays an increasing role in the study of vascular inflammatory responses. The CXCL4-induced monocyte/macrophage phenotype has previously been implicated with atherosclerotic plaque destabilization, a key process preceding plaque rupture. Monocyte-derived macrophages were found to exhibit a unique transcriptome in the presence of CXCL4 characterized by upregulation of S100A8 and MMP7. However, the mechanisms involved in CXCL4-induced monocyte-mediated vascular inflammation are unknown.

**Methods:** Single-cell RNA sequencing data were examined for CXCL4-dependent gene expression signatures in plaque macrophages. Human PBMCs were differentiated with CXCL4 and subsequently characterized in terms of osteogenic gene and protein expression signatures and calcific extracellular vesicle release. Association of the CXCL4-induced phenotype with the Wnt pathway was investigated, and CXCL4-induced PBMC-derived EV were analyzed for their potential to elicit an inflammatory response in vSMC. In-vitro findings were verified histologically in calcified human carotid artery plaques.

**Results:** In human plaque macrophages, single-cell sequencing revealed a CXCL4-susceptible subpopulation bearing a distinct proinflammatory gene expression profile. CXCL4-differentiated PBMCs exhibited a marked induction of S100A8, MMP7 and osteogenic marker transcription concomitant with augmented release of calcific EVs enriched with MMP7, S100A8 and alkaline phosphatase. Under osteogenic conditions, increased overt calcification of the extracellular matrix was observed *in vitro*. Analysis of inflammatory pathway activation identified the paracrine Wnt5a-CaMKII signaling axis to be causally linked to the CXCL4-induced osteogenic PBMC phenotype, S100A8 and MMP7 enrichment as well as calcific potential of secreted EV. Additionally, CXCL4-polarized PBMC-derived EV differentially stimulated osteogenic/inflammatory genotype transition in vSMC. In human carotid artery plaques, occurrence of CXCL4-induced mononuclear cells coincided with Wnt5a-CaMKII pathway activation and progressive plaque calcification.

**Conclusions:** This study introduces a novel mechanism driving monocyte/macrophage-mediated extracellular matrix remodeling in calcific inflammatory responses through Wnt5a-CaMKII-activated secretion of MMP7^+^S100A8^+^ calcifying EV by CXCL4-induced pro-inflammatory monocytes.

**Graphical Abstract:** 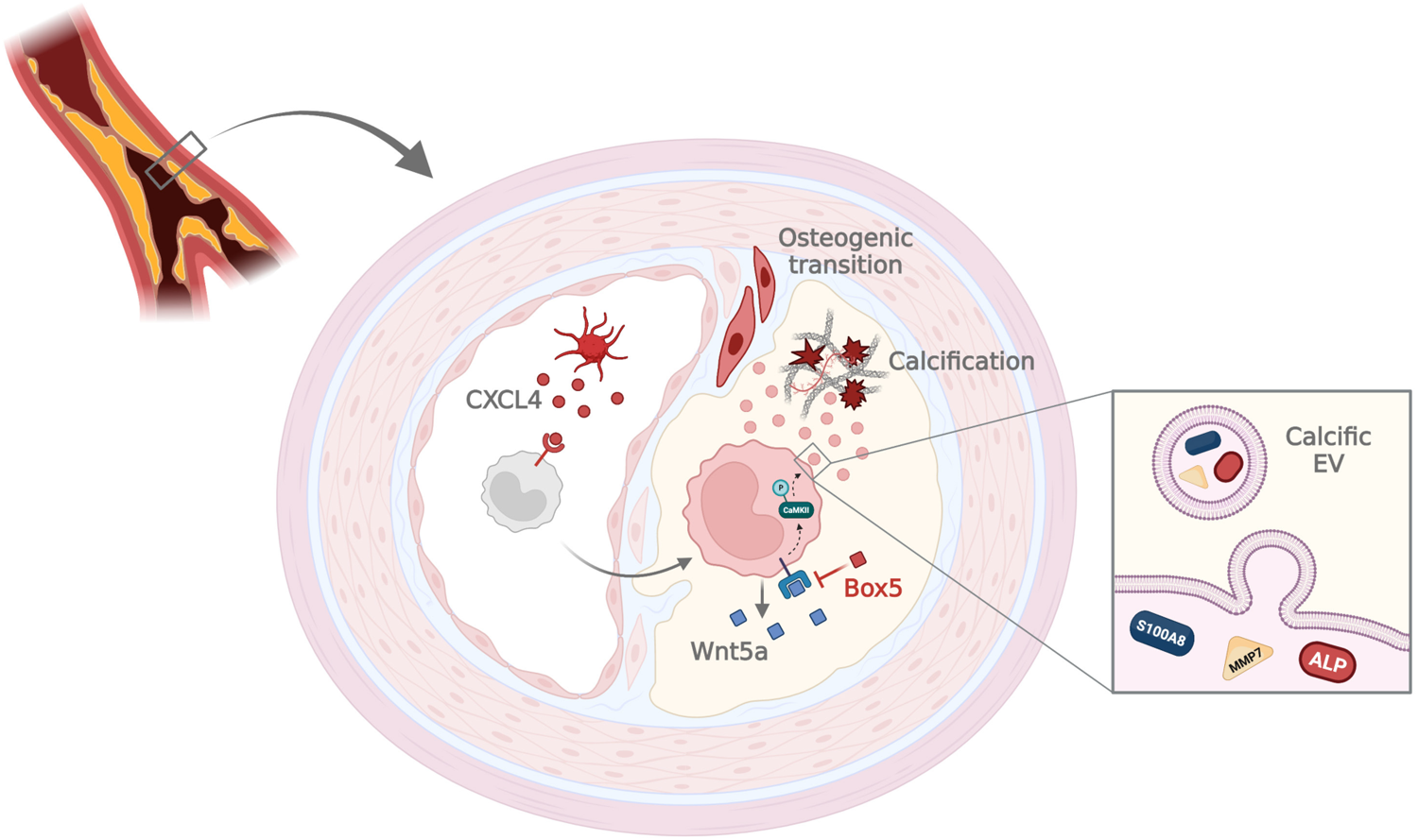

## Introduction

Cardiovascular diseases are the leading cause of death worldwide, and atherosclerosis in particular with its potentially fatal sequelae myocardial infarction and stroke account for the majority of health lost^1^. The pathophysiology of atherosclerosis has long been recognized as an inflammatory process involving effector cells pertinent to innate and adaptive immunity alike. Herein, migratory monocytes and resident macrophages have been implicated in both early and advanced stages of atheroma formation in human arterial specimen as well as in multiple experimental models of atherosclerosis^2^. Beyond the classical proinflammatory phagocyte, a phenotypic spectrum within the monocyte/macrophage lineage resulting from divergent microenvironments has been identified within the plaque, where the distinct subtypes exert pleiotropic effects in plaque genesis and growth through phenotype-specific functions^3^. The specific plaque milieu as a major determinant of morphology and resultant structural stability was found to be affected not only by vessel wall-resident cells, but also by activated platelets, whose secretome prompts a multivariate crosstalk with both resident and migratory mononuclear cell populations^4^. Systematic analysis of platelet-dependent monocyte/macrophage polarization revealed platelet-derived chemokine CXC-ligand 4 (CXCL4), also known as platelet factor 4, as a potent inducer of a unique transcriptome in monocyte-derived macrophages^5^. The specific CXCL4-invoked genotype distinguished by a marked overexpression of matrix metalloproteinase-7 (MMP7) and calcium-binding protein S100A8 has been proposed to propagate atherosclerotic plaque development, and both MMP7 and S100A8 have been independently linked to plaque progression in human tissue specimen as well as in murine atherosclerosis disease models^6, 7^. Despite propitious evidence pointing towards an unfavorable role of CXCL4-induced monocytes/macrophages in atherosclerosis, further data warranting a mechanistic integration of CXCL4-dependent inflammatory pathways on vessel cell/tissue level remain outstanding.

Extracellular vesicles (EV), lipid bilayer-enclosed nanoparticles secreted by a multitude of different cell types in various disease contexts, have been ascribed a paramount role in vascular inflammation and calcification. In the nascent plaque, EV were found to bear the potential to nucleate calcium phosphate mineral, thus budding calcific deposits of differential morphology in the vascular extracellular matrix^8^. Prior evidence demonstrated that EV secreted by pro-inflammatory macrophages exhibit calcific potential through specific enrichment of hydroxyl apatite and S100A9, thus contributing to calcification of both human and murine atherosclerotic plaques^9^.

The present study investigates the CXCL4-induced phenotype in peripheral blood-derived monocytes with respect to its potential to modulate vascular extracellular matrix under inflammatory conditions through MMP7 and S100A8 expression and calcifying extracellular vesicle release. Furthermore, our study aims to elucidate the intracellular signaling pathways conducive to CXCL4-induced monocyte polarization as potential therapeutic targets to prevent atherosclerotic plaque destabilization.

## Materials and Methods

A detailed Methods section including a Major Resources Table is available in the Data Supplement. Supporting data is available from the corresponding author upon reasonable request.

### PBMC isolation and cell culture

Fresh blood samples were obtained from healthy donors of both sexes following approval by the Institutional Review Board at the Faculty of Medicine at Heidelberg University. Isolation of peripheral blood-derived monocytes (PBMCs) was performed according to an established protocol^10, 11^. Either 100 ng/ml recombinant human M-CSF as control or 1 μM recombinant human CXCL4 were added to serum-free differentiation medium for a total of 6 days in culture. For paracrine Wnt5a stimulation, recombinant human Wnt5a (250 ng/ml) or rhWnt5a + Wnt5a-specific antagonist Box5^12^ was added for 24 hours before harvest.

### EV isolation

Extracellular vesicles were isolated from cell culture media as described previously^13–15^. Pellets were either analyzed immediately or stored at −80°C until further analysis.

### EV-vSMC coculture

EV isolates from differentially stimulated PBMCs were purified by centrifugal filtration. Equal amounts of EV were used for coculture experiments. Human vascular smooth muscle cells (vSMC) were purchased from PromoCell. VSMC between passages 5 and 9 were used for coculture experiments. After 48 hours in culture, vSMC were serum-starved and incubated with PBMC-derived EV for 24 hours before harvest.

### EV fluorescent labeling

PBMC-derived EV were fluorescently labeled using lipophilic dye PKH67 essentially as described previously^16^. Uptake of fluorescent EV by vSMC was detected using an Axio Observer Z1 fluorescence microscope (Zeiss, Oberkochen, Germany).

### Human carotid artery tissue

Frozen carotid artery punch specimens (n=10) from patients undergoing carotid thromboendarterectomy surgery at University Hospital Heidelberg were obtained from the Vascular Biomaterial Bank Heidelberg (VBBH) following Institutional Review Board approval. Besides age (73.3 ±8.8 years) and sex (90% male), no additional demographic or clinical information was made available to the authors prior to analysis.

### Immunofluorescence of carotid artery specimen

Frozen tissue sections were thawed, fixed in 4% paraformaldehyde and subsequently hydrated. Following unspecific protein blocking with 5% BSA in PBS, sections were incubated with primary antibody or concentration- and species-matched isotype control overnight. Biotinylated secondary antibody was added for one hour, followed by fluorophore-conjugated streptavidin for 30 mins. For double/triple immunofluorescence, all steps starting with primary antibody were repeated once/twice, each time preceded by an avidin/biotin blocking step to avoid fluorescent labeling of unbound biotin from the previous step. Images were obtained on a Leica SP8 confocal microscope. For each tissue section, positive cells were counted using the Leica Application Suite X software version 3.5.7, and the number of positive cells was divided by total tissue area.

### Single-cell RNA sequencing analysis

A single-cell sequencing database of human coronary plaque tissue was accessed through the Gene Expression Omnibus platform (accession code GSE131778). Analysis was conducted using an established protocol^17^ with R package Seurat version 4.3.0. Of the resulting cell clusters, a subset containing monocyte-derived macrophages and resident macrophages was isolated and cluster identity was reassigned based on dichotomized *CCR1* expression readout. Differentially expressed genes in *CCR1^+^* plaque mononuclear cells and respective Gene Ontology (GO) annotations were identified by enrichment analysis using the Enrichr tool as previously described^18^.

## Results

### Single-cell RNA sequencing of human coronary plaque tissue identifies a distinct gene expression signature in CXCL4-susceptible macrophages

Previous gene expression analyses have ascribed a distinct genotype to peripheral blood-derived monocytes (PBMC) differentiated in the presence of CXCL4 *in vitro*, bearing a gene expression profile divergent from those found in established monocyte/macrophage subtypes^5^. In an effort to verify the existence of a CXCL4-susceptible mononuclear cell population with specific genotypic features suggestive of its functional role in the developed atherosclerotic plaque, a single-cell RNA sequencing repository of coronary atheromata from explanted human hearts was examined^17^ (Fig. 1A). Among monocyte-derived and resident macrophage clusters characterized by expression of cell adhesion-associated macrosialin CD68, approximately 28% of cells were found to express CC-chemokine receptor 1 (CCR1), which has previously been identified as the primary receptor of CXCL4 ligand interaction in mononuclear cells^19^ (Fig. 1B). Differentially expressed genes in CCR1^+^ cells of the plaque macrophage cluster encompassed RNA transcripts for calcium-binding complexes calprotectin (encoded by S100A8 and S100A9 genes) and calgranulin C (encoded by S100A12) as well as GCA coding for the grancalcin protein (Fig. 1C+D). Functional enrichment analysis exploiting the Gene Ontology (GO) resource found a predisposition of target proteins of differentially expressed CCR1^+^ macrophage RNA transcripts in the secretory granule/vesicle lumen and membrane compartments (Fig. 1E, S1A). Predominant GO Molecular Function annotations included chemokine-associated activity and cytokine-related receptor interactions (Fig. 1F, S1B). Conclusively, plaque-derived single-cell RNA sequencing data support a subpopulation of CXCL4-susceptible macrophages presuming a proinflammatory gene expression signature selectively targeting the secretory vesicle membrane and lumen compartment.

**Figure 1:**
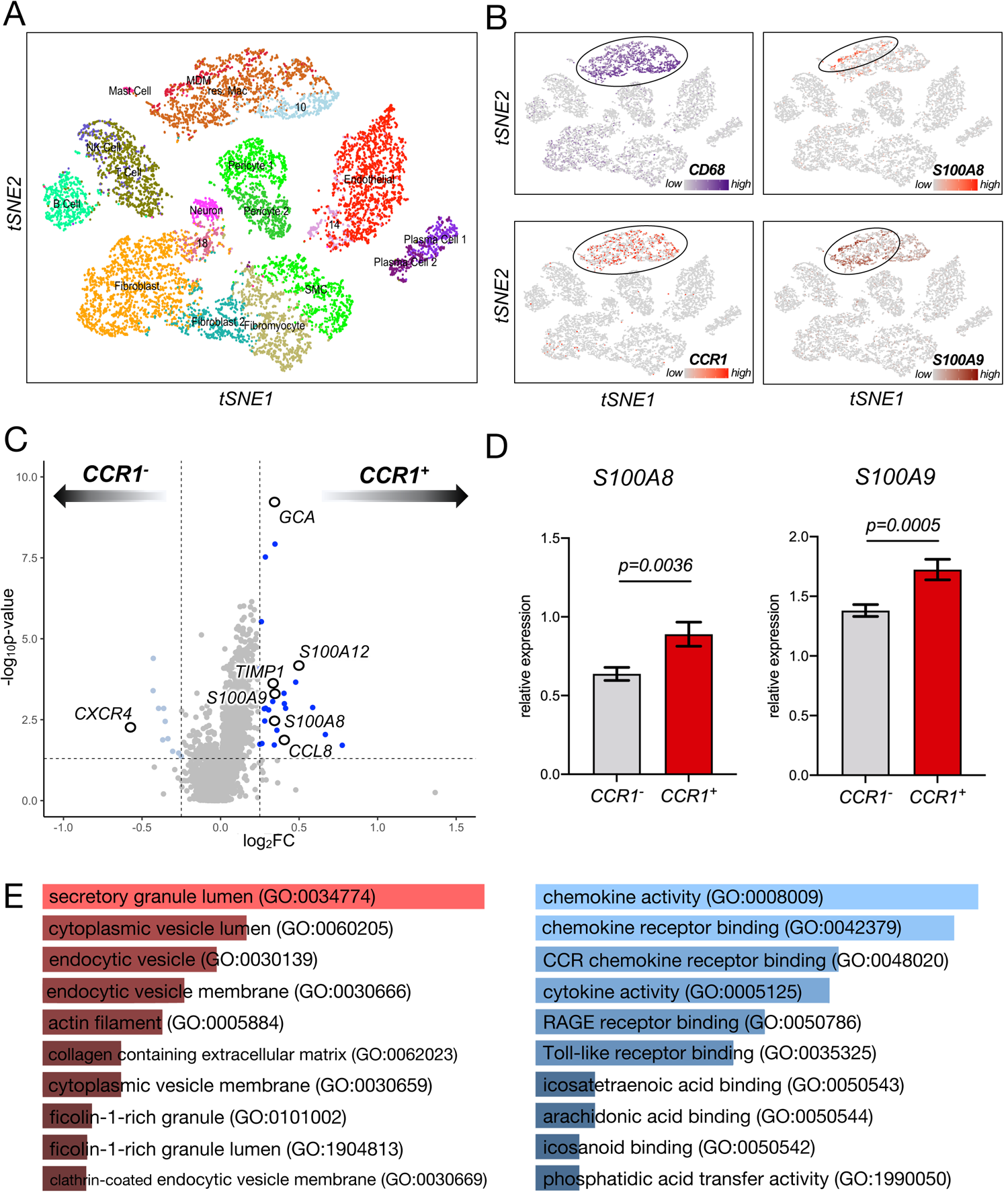
Single-cell RNA sequencing of human coronary artery plaque tissue identifies a population of CXCL4-susceptible macrophages with distinct gene expression profile. **A** t-SNE visualization of cell clusters in right coronary artery plaques from n=4 patients. Cluster numbers represent indiscriminately defined cell clusters. *MDM: monocyte-derived macrophages*, *res. Mac: resident macrophages, SMC: smooth muscle cells.* **B** t-SNE cluster visualization with gene expression overlay of CD68 (*top left*), CCR1 (*bottom left*), S100A8 (*top right*), and S100A9 (*bottom right*). **C** Volcano Plot of differentially expressed genes between CCR1^+^ and CCR1^-^ plaque mononuclear cells, significantly upregulated genes are highlighted in dark blue for CCR1^+^ and in light blue for CCR1^-^ cells. Genes of interest among differentially regulated genes are highlighted and labeled. **D** Mean relative expression of S100A8 (*left*) and S100A9 (*right*) in CCR1^+^ vs. CCR1^-^ plaque mononuclear cells. *Data shown as mean±SEM, n=1185 CCR1^-^ vs. 458 CCR1^+^ cells from n=4 biological replicates. Wilcoxon’s rank sum test; genes with associated p-value <0.05 and log_2_FC threshold of >0.25 for CCR1^+^ and <-0.25 for CCR1^-^ were considered differentially expressed.* Gene Ontology (GO) terms in Cellular Component (**E**) and Molecular Function (**F**) annotations associated with top 100 differentially expressed genes in CCR1^+^ plaque mononuclear cells identified by functional enrichment analysis. Terms ranked by p-value; specific GO terms in parentheses. *Fisher’s exact test with Benjamini-Hochberg adjustment for multiple testing*.

### CXCL4 induces distinct gene and protein expression signatures in PBMCs *in vitro*

To further elucidate the mechanisms of CXCL4-induced monocyte maturation, CXCL4-specific gene and protein expression signatures were evaluated *in vitro*. Compared to M-CSF maturation, CXCL4 stimulation of naïve PBMCs *in vitro* resulted in a specific genotype characterized by upregulated mRNA expression of MMP7 and calcium-binding protein S100A8, confirming previous analyses^5, 20^ (Fig. 2A). Further, a significant upregulation of osteogenic markers ALP transcribing the alkaline phosphatase protein, and OPN encoding calcium-binding sialoprotein osteopontin is appreciated (Fig. 2B). Conversely, differentiation by CXCL4 versus M-CSF triggers downregulation of pattern recognition co-receptor CD14 and Fcγ receptor CD16 on gene expression level (Fig. 2A) as well as downregulation of CD68 on the surface of differentially stimulated PBMCs (Fig. 3B). On protein expression level, a significant induction of S100A8 and annexin 5, which has previously been found to colocalize with S100 family members to facilitate mineralization^21^, along with upregulated MMP7 protein expression (Fig. S2, S3) was observed in CXCL4-differentiated PBMCs compared to M-CSF control (Fig. 2D). Thus, *in vitro* differentiation of PBMCs with CXCL4 favors multiple gene and protein markers pertinent to a pro-calcific and/or pro-osteogenic phenotype.

**Figure 2:**
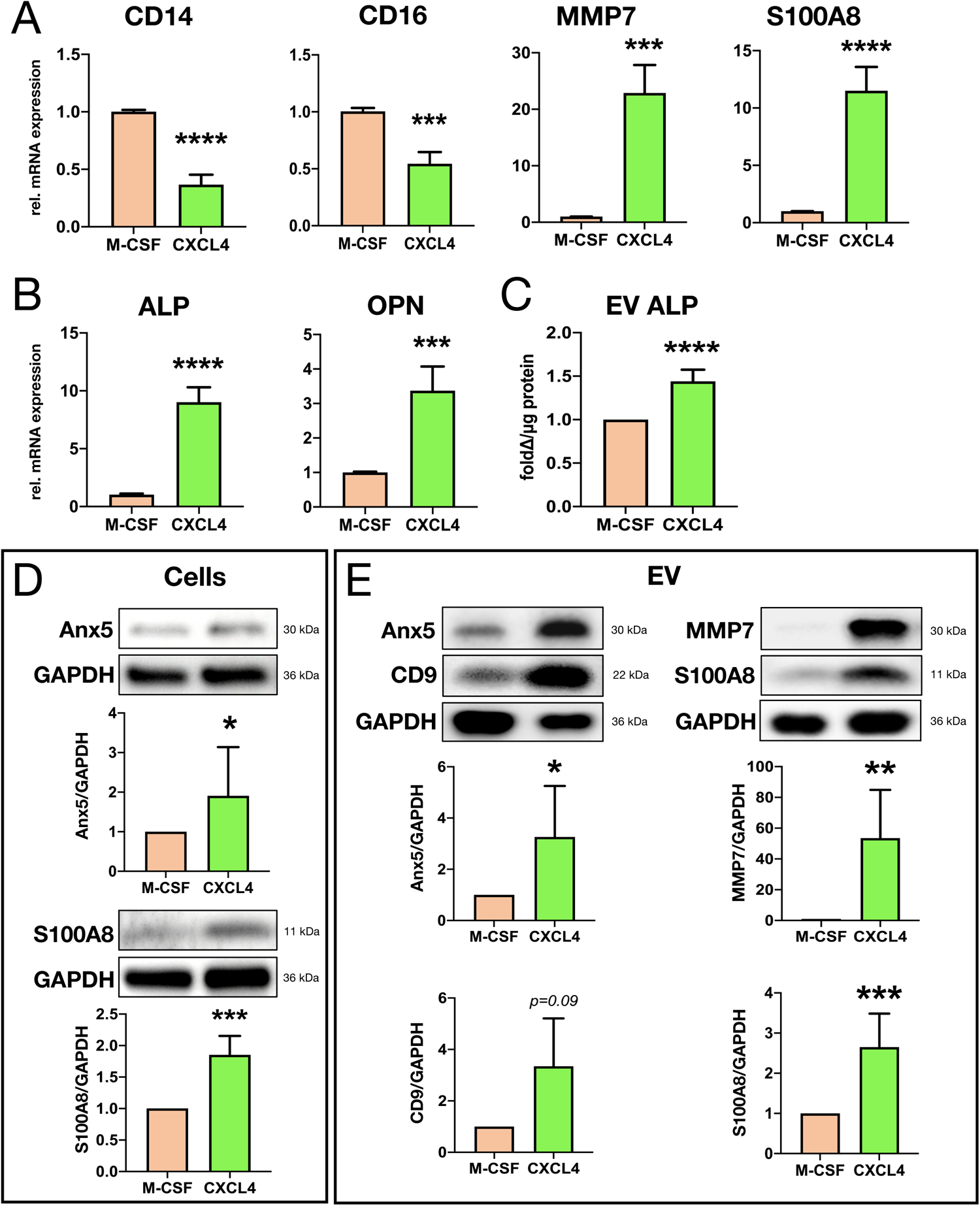
CXCL4 induces the release of extracellular vesicles (EV) and downregulation of macrosialin CD68 compared to M-CSF-differentiated PBMCs. **A** Representative brightfield images at 40x magnification of human PBMCs differentiated with M-CSF (*top*) or CXCL4 (*bottom*) after 6 days in culture. *n=8 biological replicates, scale bar 50 μm.* B Immunofluorescence for CD68 expression in M-CSF-(*top panel*) or CXCL4-induced PBMCs (*center panel*) after 6d in culture at 63x magnification; *right*: quantification of mean fluorescence intensity (MFI) compared to secondary antibody-only control. *Data shown as mean±SD, n=5 biological replicates per group. Shapiro-Wilk normality test followed by one-way ANOVA with Bonferroni post-hoc test;* ^##^*p<0.01 ****p<0.0001*. **C** Particle size distribution analyzed by nanoparticle tracking analysis (NTA) of EV isolated from cell culture media of M-CSF-(*top*) or CXCL4-induced PBMCs (*bottom*) after 24h in culture. *Mean±SEM at 20 nm increments, n=4-6 biological replicates per group*. **D** Mean particle size (*top*) and particle concentration (*bottom*) of EV isolated from differentiated PBMCs in culture, quantified by NTA. *n=4-6 biological replicates, Shapiro-Wilk normality test followed by mixed-effects analysis with Bonferroni post-hoc test; *p<0.05*.

**Figure 3:**
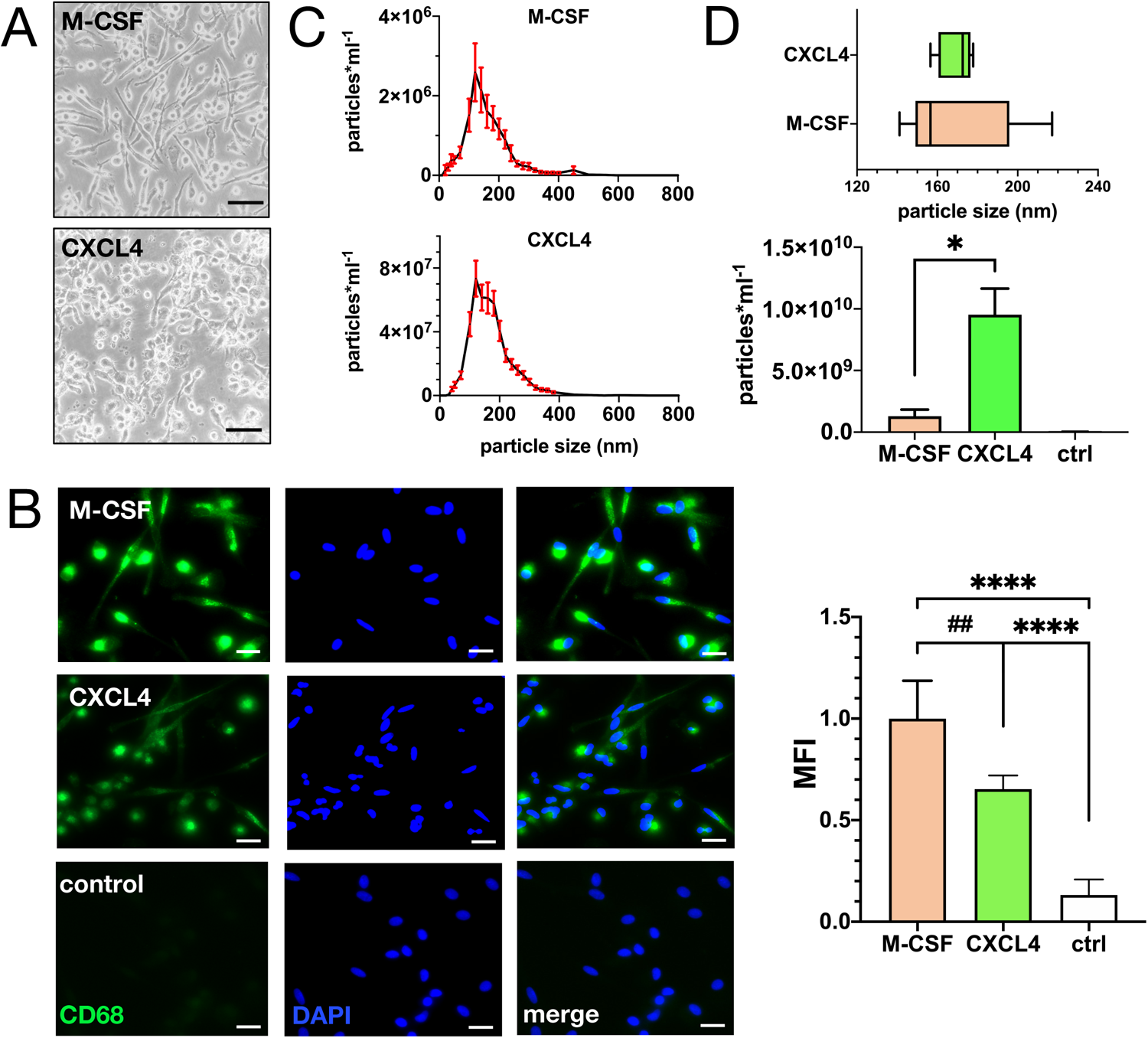
CXCL4 propagates a distinct gene expression signature in cultured PBMCs and a specific protein cargo in PBMC-derived EV. **A** Relative mRNA expression levels of CD14, CD16, MMP7 and S100A8 and **B** osteogenic markers alkaline phosphatase (ALP) and osteopontin (OPN) following CXCL4 induction, normalized to M-CSF. *Mean±SEM, Shapiro-Wilk normality test followed by either unpaired student’s t-test or Mann-Whitney U for non-normally distributed parameters, n=4-6 biological replicates per group.* **C** ALP activity in EV derived from CXCL4-induced PBMC normalized to M-CSF. *Mean±SD, Shapiro-Wilk normality test followed by unpaired student’s t-test, n=6 biological replicates*. **D** Representative immunoblots for annexin 5 (Anx5; *top*) and S100A8 (*bottom*) from cell lysates of CXCL4- vs. M-CSF-induced PBMCs. *Mean±SD, Shapiro-Wilk normality test followed by either unpaired student’s t-test or Mann-Whitney U for non-normally distributed parameters, n=5 biological replicates.* **E** Representative immunoblots for Anx5, CD9, MMP7 and S100A8 from PBMC-derived EV isolates. *Mean±SD, Shapiro-Wilk normality test and unpaired student’s t-test, n=3-4 biological replicates. *p<0.05 **p<0.01 ***p<0.001 ****p<0.0001*.

### CXCL4 stimulation of PBMCs facilitate the release of calcific EV loaded with MMP7 and S100A8 *in vitro*

Single-cell RNA sequencing analysis and present *in vitro* data equally suggest osteogenic and EV-enriched expression signatures in CXCL4-polarized monocytes/macrophages, proposing a potential implication of CXCL4 stimulation on EV release in mononuclear cells. To test this hypothesis, nanoparticle tracking of EV isolated from CXCL4-versus M-CSF-differentiated PBMCs was performed. Herein, EV populations of typical size distribution and similar average particle size (*p=0.85*; Fig. 3C, D) were visualized. Quantitative analysis of CXCL4-compared to M-CSF-induced PBMC-derived EV showed a ∼7-fold higher concentration of EV released in vitro under baseline conditions (9.53e9 ± 2.12e9 times ml^-1^ for CXCL4 vs. 1.29e9 ± 5.38e8 times ml^-1^ for M-CSF, *p=0.03*; Fig. 3D). Further evaluation of EV protein content yielded an approximately 44% increase in relative alkaline phosphatase activity in EV isolates from CXCL4- vs. M-CSF-induced PBMCs (*p<0.0001*; Fig. 2C), and CXCL4 differentiation resulted in higher expression of potential mineralization facilitators Annexin 5 and S100A8, tetraspanin CD9 as well as a marked elevation of MMP7 in PBMC-derived EV (Fig. 2E). Hence, CXCL4 differentiation of PBMCs stimulated EV release and EV calcific potential as well as MMP7 content.

### Osteogenic conditions favor CXCL4-induced MMP7 and S100A8 expression and activate noncanonical Wnt5a-mediated signaling in PBMCs

Osteogenic culture conditions mimicking local plaque milieu were commonly found to aggravate pathogenic expression profiles in susceptible cell populations. Supplementation of β-glycerophosphate, a synthetic substrate of alkaline phosphatase, and calcium to PBMC differentiation media to facilitate an osteogenic culture milieu invoked a synergistic effect on the CXCL4-induced genotype: Osteogenic conditions markedly bolstered CXCL4-mediated overexpression of MMP7, S100A8 and OPN gene transcription whilst not impacting upregulation of ALP mRNA in CXCL4-differentiated PBMCs *in vitro* (Fig. 4B). In EV, however, a significant increase of ALP activity under osteogenic conditions was observed (Fig. 4C), concomitant with augmented Alizarin Red staining intensity of cultured PBMCs as a hallmark of overt extracellular matrix calcification (Fig. 4A). These data suggest a specific enhancement of ALP loading into EV in an osteogenic environment.

**Figure 4:**
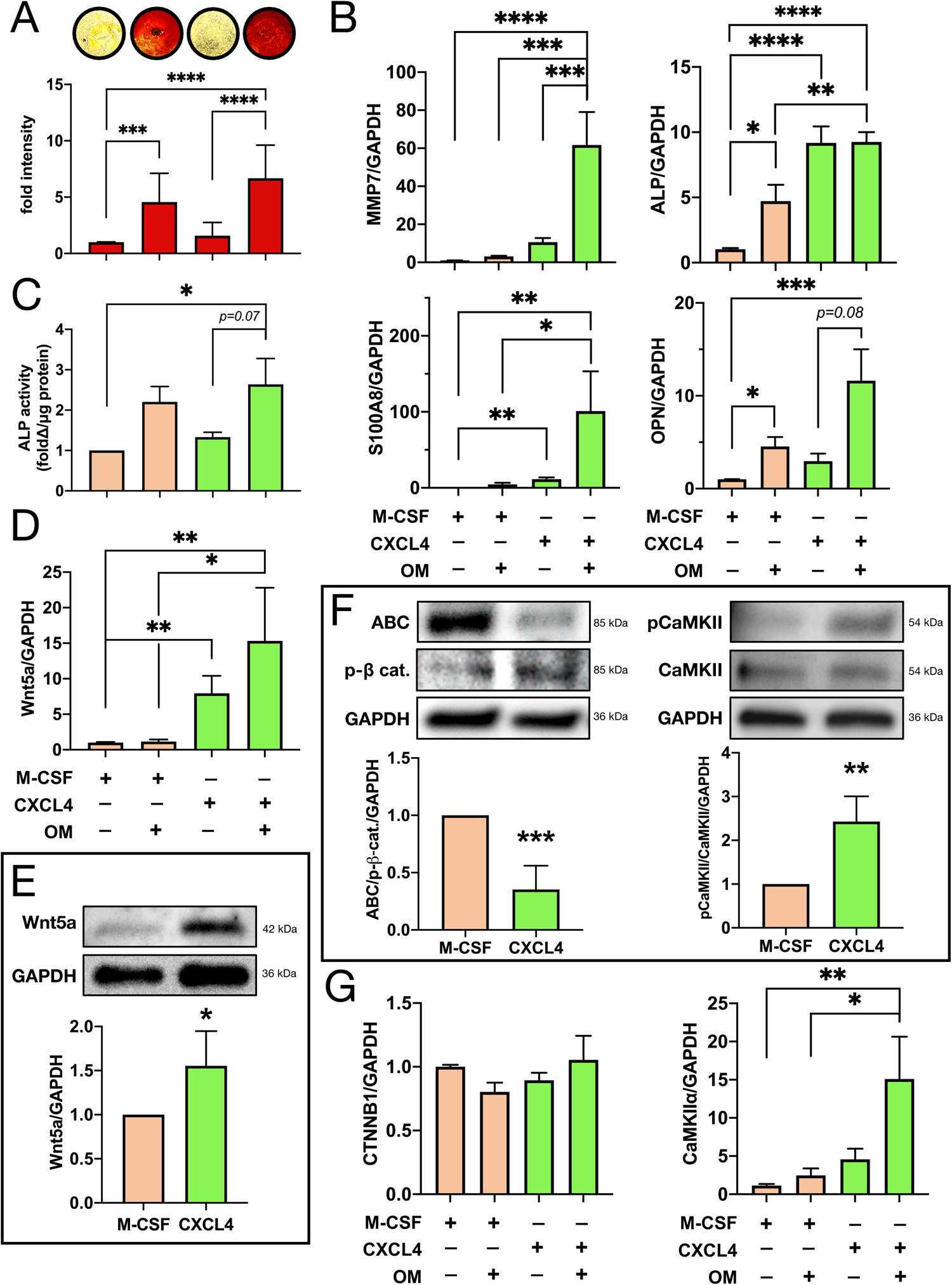
CXCL4-differentiated PBMCs exhibit augmented overt calcification potential concomitant with increased osteogenic gene expression and activation of non-canonical Wnt5a-Ca^2+^ signaling under osteogenic conditions *in vitro*. **A** Brightfield images at 4x magnification (*top panel*) and quantification (*bottom*) of Alizarin Red S staining of M-CSF- vs. CXCL4-induced PBMCs cultured in the presence or absence of osteogenic medium (OM). *Mean±SD, Shapiro-Wilk normality test and Kruskal-Wallis test with Dunn’s multiple comparisons test, n=8 biological replicates.* B Relative mRNA expression of MMP7, S100A8, ALP, OPN and **D** Wnt5a after differentiation with either M-CSF or CXCL4 and in the presence or absence of osteogenic media. *Mean±SEM, Shapiro-Wilk normality test and either mixed-effects analysis with Bonferroni post-hoc test or Kruskal-Wallis test with Dunn’s multiple comparisons test for non-normally distributed parameters, n=3-4 biological replicates*. **C** ALP activity in EV from PBMCs cultured with M-CSF or CXCL4 with or without osteogenic media. *Mean±SEM, Shapiro-Wilk normality test and mixed-effects analysis with Bonferroni post-hoc test, n=3-6 biological replicates.* E Representative immunoblots and quantification of Wnt5a protein, **F** active β-catenin and phosphorylated Calcium-Calmodulin Kinase II (CaMKII) under CXCL4 or M-CSF stimulation. *Mean±SD, Shapiro-Wilk normality test and unpaired student’s t-test, n=4-5 biological replicates.* G Relative mRNA expression of CTNNB1 and CaMKIIα with either M-CSF or CXCL4 differentiation and osteogenic media. *Mean±SEM, Shapiro-Wilk normality test and either mixed-effects analysis with Bonferroni post-hoc test or Kruskal-Wallis test with Dunn’s multiple comparisons test for non-normally distributed parameters, n=3-5 biological replicates*. *\*p<0.05 **p<0.01 ***p<0.001 ****p<0.0001*.

Several contributors to the Wnt pathway have previously been implicated in the expression of both MMP7 and S100A8 in monocyte/macrophage-mediated extracellular matrix modulation in various disease contexts^22, 23^. Relative gene expression for common Wnt targets was investigated in PBMCs differentiated with M-CSF, CXCL4 or a combination of interferon γ (IFNγ) and lipopolysaccharide (LPS) to induce a conventional monocyte proinflammatory genotype. Among differentially regulated Wnt genes, expression levels of Wnt5a in particular exhibited pronounced upregulation in CXCL4- and IFNγ+LPS-induced PBMCs (Fig. S4, Supp. Table 1), of which the latter has previously been ascribed a crucial role in the macrophage-mediated inflammatory response^24^. Upregulation of Wnt5a on gene expression level was further augmented in the presence of osteogenic media in CXCL4-, but not in M-CSF-differentiated PBMCs (Fig. 4D), and a concomitant induction of Wnt5a protein on cell level was appreciated in the presence of CXCL4 (Fig. 4E).

In previous studies, Wnt5a has been associated with several noncanonical signaling pathways^25^. Concordantly, CXCL4 differentiation of PBMCs suppressed active relative to phosphorylated beta catenin, a key factor in canonical Wnt signaling, compared to M-CSF control (Fig. 4F), while no difference in mRNA expression of target gene CTNNB1 was found in any condition (Fig. 4G). Further analysis of common noncanonical downstream signaling targets revealed a significant upregulation of calcium calmodulin kinase II (CaMKII) phosphorylation in CXCL4-compared to M-CSF-induced PBMCs (Fig. 4F) with concurrent incremental expression of CaMKII α-subunit mRNA in the presence of CXCL4 and osteogenic conditions (Fig. 4G). Further investigation into common noncanonical downstream regulators showed consistent reciprocal downregulation of the phosphorylation of Erk MAP kinase (Fig. S5A), c-Jun N-terminal kinase (JNK; Fig. S5B) or p38 (Fig. S5C) in PBMCs following CXCL4 differentiation, underlining presented evidence of a specific Wnt5a-pCaMKII-mediated signaling activation conducive to the CXCL4-induced phenotype.

### The CXCL4-induced PBMC phenotype and associated calcific EV release exhibit auto-/paracrine Wnt5a dependency

The proinflammatory macrophage phenotype has been causatively linked to auto-/paracrine Wnt5a ligand/receptor interaction in a previous study^24^. Dependency of the CXCL4-differentiated PBMC phenotype on Wnt5a-mediated noncanonical signaling activation was investigated by recombinant Wnt5a stimulation of CXCL4-differentiated PBMCs and reciprocal inhibition of Wnt5a-specific receptor activation by Wnt5a hexapeptide analogue Box5^12^ (Fig. 5A). Exogenous Wnt5a stimulation markedly augmented quantitative EV release *in vitro* in CXCL4-differentiated PBMCs by about ten-fold (4.43e10 ± 6.38e9 times ml^-1^ in CXCL4/Wnt5a vs. 4.03e9 ± 7.80e8 times ml^-1^ in CXCL4; *p<0.0001*; Fig. 5B), concomitant with a significant induction of EV calcific potential measured by relative ALP activity in EV (Fig. 5B) as well as EV loading with Annexin 5, CD9, S100A8 and MMP7 (Fig. 5F), all of which cumulatively resulted in enhanced overt calcification by Wnt5a-stimulated CXCL4-differentiated PBMCs *in vitro* (Fig. 5C) compared to M-CSF control. Inhibition of Wnt5a ligand-receptor interaction by Box5 consistently reversed all aspects of the Wnt5a-induced phenotype in CXCL4-differentiated PBMCs including quantitative EV release and EV calcific potential (Fig. 5B), relative expression of Annexin 5, CD9, S100A8 and MMP7 in EV (Fig. 5F), as well as Alizarin Red S-positive extracellular matrix calcification (Fig. 5C).

**Figure 5:**
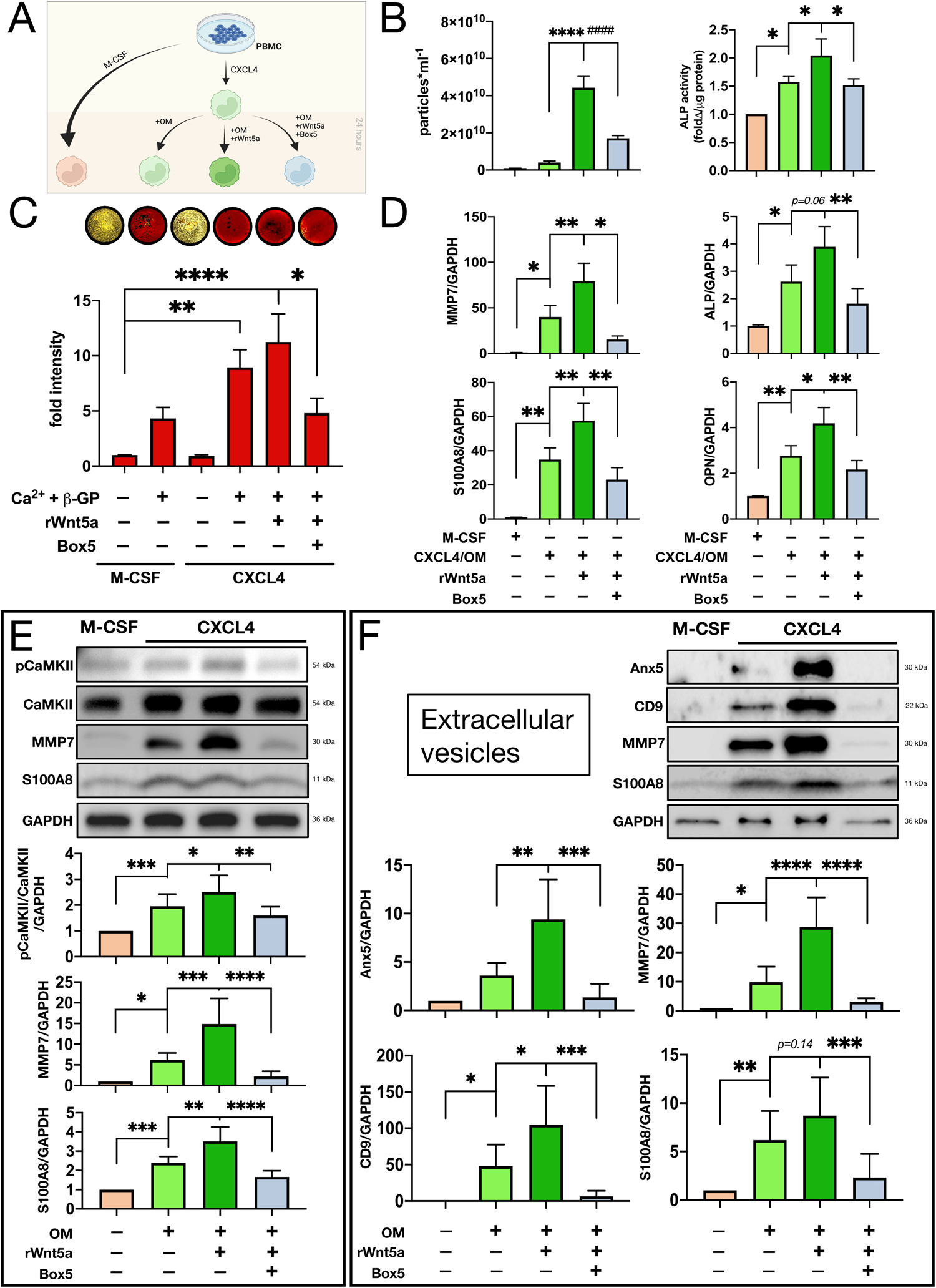
Stimulation of CXCL4-induced PBMCs with exogenous Wnt5a and reciprocal Wnt5a-specific inhibition with small peptide inhibitor Box5 shows Wnt5a dependency of EV-mediated calcification potential, osteogenic genotype as well as MMP7 expression and EV loading. **A** Schematic of differential PBMC stimulation (*created with BioRender*). **B** EV concentration determined by nanoparticle tracking and ALP activity of EV isolates from differentially stimulated PBMCs. *Mean±SEM, Shapiro-Wilk normality test and mixed-effects analysis with Bonferroni post-hoc test, n=3-6 biological replicates*. **C** Brightfield images (*top panel*) of Alizarin Red S stained PBMCs following differential stimulation and staining quantification (*bottom*). *Mean±SD, Shapiro-Wilk normality test and Friedman test with Dunn’s multiple comparison test, n=3 biological replicates.* D Relative mRNA expression of MMP7, S100A8, ALP and OPN in differentially stimulated PBMCs. *Mean±SEM, mixed-effects analysis with Bonferroni post-hoc test, n=3-6 biological replicates.* E Representative immunoblots and quantification for phospho-CaMKII, MMP7 and S100A8 from cell lysates of differentially stimulated PBMCs. *Mean±SD, Shapiro-Wilk normality test and mixed-effects analysis with Bonferroni post-hoc test, n=4-7 biological replicates.* F Representative immunoblots and quantification for Anx5, CD9, MMP7 and S100A8 from EV isolates of differentially stimulated PBMCs. *Mean±SD, Shapiro-Wilk normality test and One-way ANOVA with Bonferroni post-hoc test, n=4 biological replicates. *p<0.05 **p<0.01 ***p<0.001 ****^/####^p<0.0001*.

On cell level, congruent effects of exogenous Wnt5a that were attenuated by addition of Box5 were observed for the CXCL4-induced pro-osteogenic genotype characterized by mRNA expression of MMP7, S100A8, ALP and OPN (Fig. 5D), and the CXCL4-mediated protein expression signature encompassing MMP7 and S100A8 as well as CaMKII activation (Fig. 5E).

### EV released by CXCL4-induced PBMCs promote a pro-calcific, pro-inflammatory genotype in vascular smooth muscle cells

EV as mediators of cell-cell communication have been identified as a driving force in inflammatory processes of the vasculature^26^, and EV derived from migratory leucocytes in particular were found to impact resident vascular cells to propagate inflammatory responses^27^. To investigate a putative effect of EV released by PBMCs differentiated with CXCL4 alone or in combination with Wnt5a on the vascular smooth muscle cell (vSMC) genotype, EV isolated from differentiated PBMCs in culture were incubated with naïve human vSMC (Fig. 6A). Intracellular uptake of isolated PBMC-derived EV by vSMC *in vitro* was visualized by fluorescent EV labeling (Fig. 6B). Compared to vSMC grown in control media, under osteogenic conditions, or with addition of M-CSF-induced PBMC-derived EV, co-culture with EV isolated from PBMCs stimulated by either CXCL4 alone or CXCL4 combined with Wnt5a led to incremental induction of alkaline phosphatase gene expression in naïve vSMC (Fig. 6C), indicating osteogenic genotype transition. For interleukin 6, a key marker of cellular inflammatory responses, a significant upregulation of gene expression in vSMC co-cultured with EV from CXCL4- and Wnt5a-stimulated PBMCs was observed compared to control vSMC as well as vSMC cultured with EV from PBMCs stimulated by M-CSF or CXCL4 alone (Fig. 6D). No difference in the expression of osteogenic gene markers osteopontin or Runx2 (Fig. S6B), nor of alpha-smooth muscle actin, collagen type I alpha-1 or fibronectin-1, previously identified gene markers of vSMC phenotypic switch, was appreciated after co-culture with CXCL4- or CXCL4-/Wnt5a-stimulated PBMC-derived EV (Fig. S6A). Thus, PBMCs stimulated with CXCL4 in combination with Wnt5a exhibited the potential to propagate a pro-inflammatory and pro-calcific state in proximal vSMCs by EV-mediated paracrine signaling.

**Figure 6:**
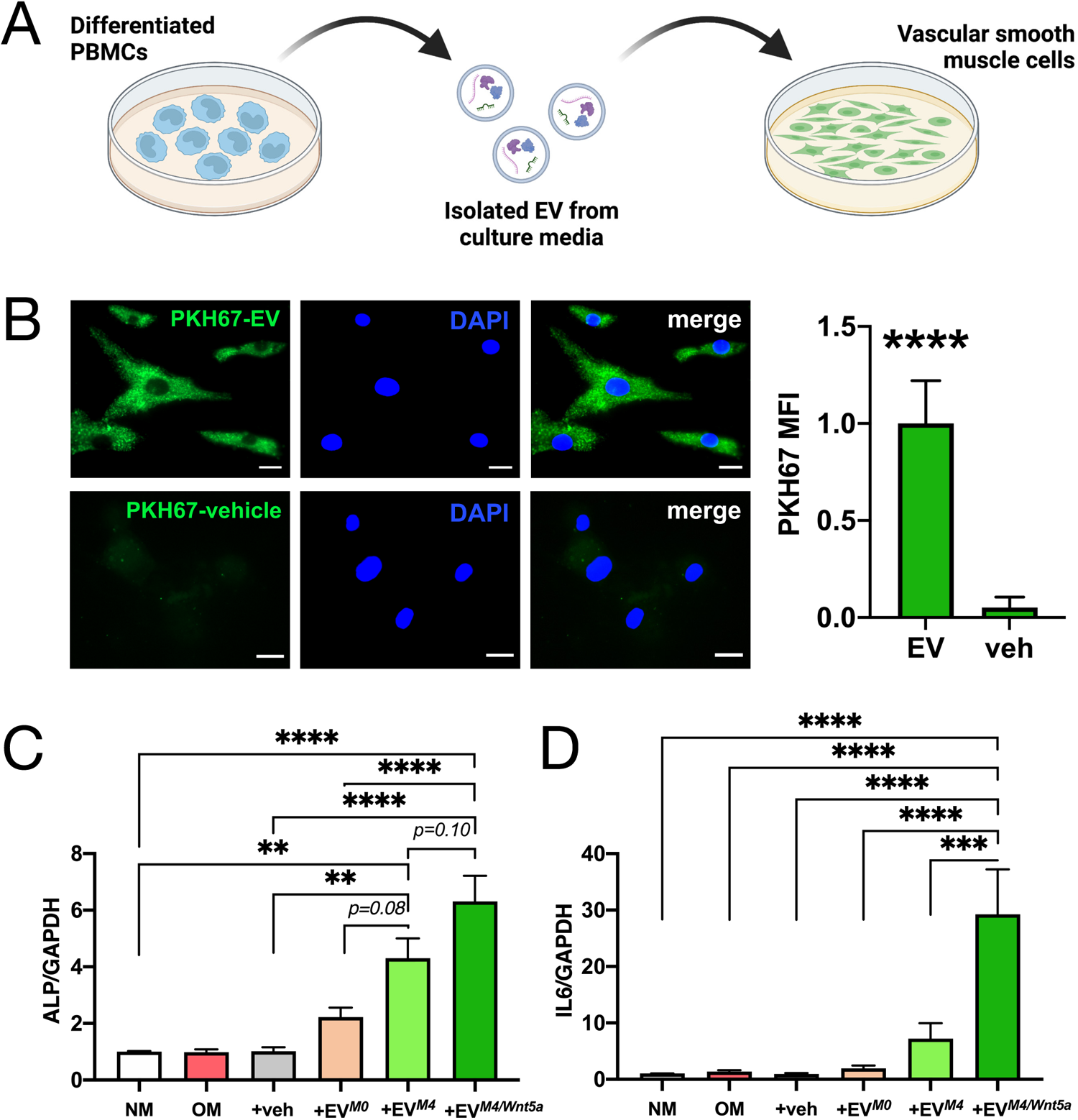
Conditioning with PBMC-derived EV induces key inflammatory and procalcific genes in vascular smooth muscle cells. **A** Schematic of PBMC-derived EV isolation and subsequent coculture with vSMC. *Created with BioRender*. **B** Representative immunofluorescent images of PKH67-labeled PBMC-derived EV (*top panel*) or vehicle (*bottom panel*) in vSMC after 24 hours of coculture. *Right*: Quantification of PKH67 mean fluorescence intensity (MFI) of labeled EV vs. vehicle (veh). *Mean±SD, Shapiro-Wilk normality test followed by unpaired student’s t-test, n=3 biological replicates. Scale bar 20 μm.* **C** Relative mRNA expression levels of ALP or **D** interleukin 6 (IL6) of vSMC conditioned with osteogenic media (OM), vehicle (veh), EV from PBMC differentiated with M-CSF (EV*^M0^*), CXCL4 (EV*^M4^*), or CXCL4 and exogenous Wnt5a (EV*^M4/Wnt5a^*), normalized to GAPDH and unstimulated vSMC control (NM). *Mean±SEM, Shapiro-Wilk normality test followed by mixed-effects analysis with Bonferroni post-hoc test, n=4 vSMC lines with n=3 PBMC donors, respectively. **p<0.01 ***p<0.001 ****p<0.0001*.

### In human *ex-vivo* carotid artery plaques, advanced lesion calcification associates with CD68^+^MMP7^+^S100A8^+^ cell abundance and Wnt5a-Ca^2+^ pathway activation

To verify our *in vitro* findings on the CXCL4-induced PBMC phenotype and its association with calcific EV release and Wnt5a-CaMKII pathway activation in an *ex vivo* disease model of atherosclerosis, extra-operative tissue samples from atherosclerotic human carotid arteries at varying disease stages were examined by immunohistochemistry. In plaques exhibiting advanced fibro-calcification (Fig. 7A), a spatial correlation of Wnt5a expression in and around the calcified core could be observed, and progressive plaque calcification positively correlated with Wnt5a immunostaining (Fig. 7B) as described in a previous study^28^. The occurrence of CXCL4-induced mononuclear cells in plaque tissue was verified by triple immunostaining for CD68, MMP7 and S100A8 as previously described^20^ (Fig. 7C,D), and incidental plaque calcification was found to significantly coincide with local CD68^+^MMP7^+^S100A8^+^ cell abundance (Pearson’s R^2^ = 0.48, p=0.001; Fig. 7F). Concordantly, phosphorylated CaMKII in CD68^+^ cells in the plaque (Fig. 7E) significantly correlated with both CD68^+^MMP7^+^S100A8^+^ cell density (Pearson’s R^2^ = 0.51, p<0.001) and overt plaque calcification (Pearson’s R^2^ = 0.67, p=0.008; Fig. 7G).

**Figure 7:**
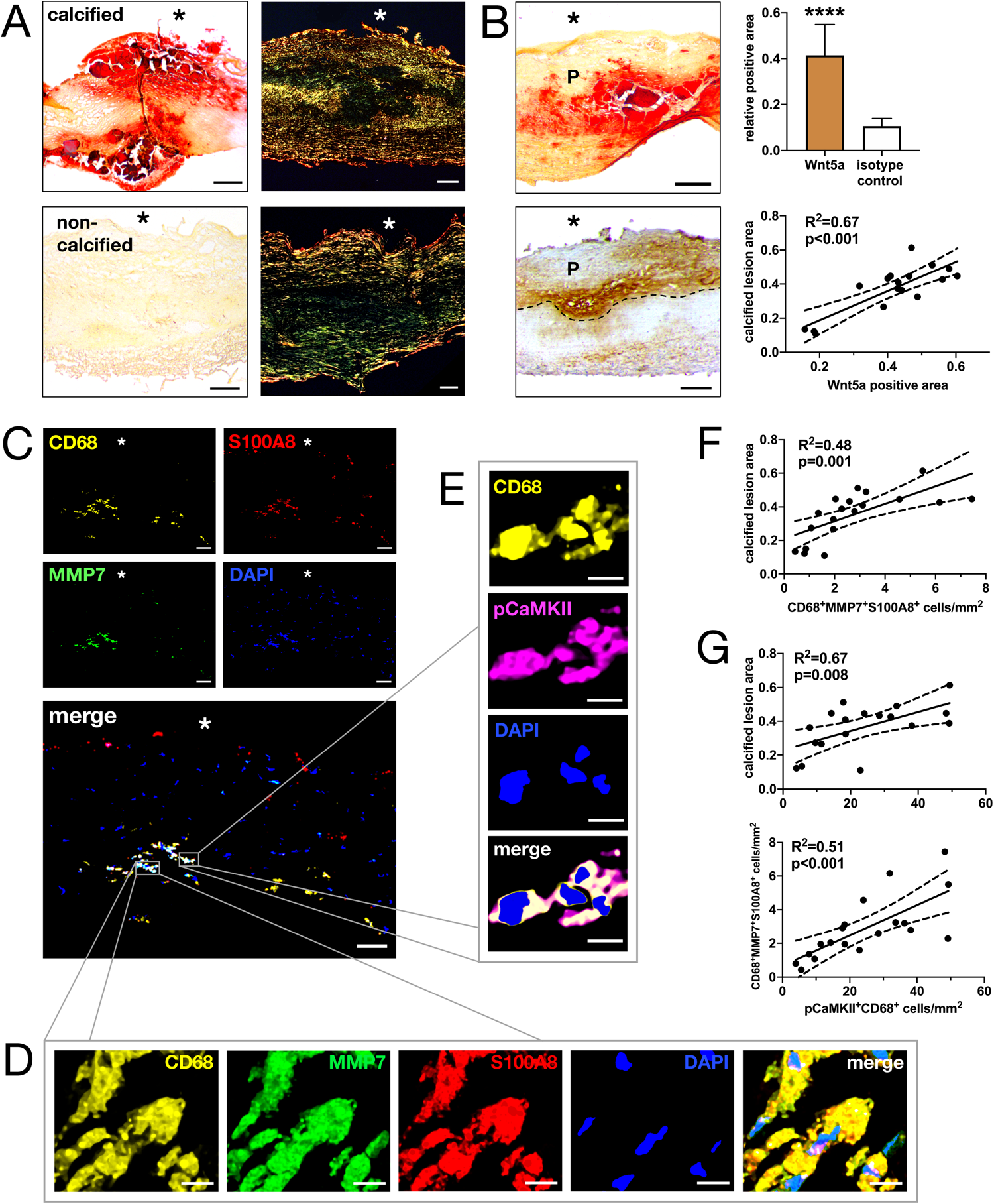
Immunohistochemical analysis of ex-vivo human carotid artery plaques recapitulates multifactorial associations between CD68^+^MMP7^+^S100A8^+^ mononuclear cell abundance, noncanonical Wnt5a-Ca^2+^ pathway activation and progressive plaque calcification. **A** Representative brightfield images of Alizarin Red S (*left*) and polarized images of Picrosirius Red staining (*right*) of calcified (*top panel*) and noncalcified (*bottom panel*) carotid artery plaques at 4x magnification. *Scale bar 200 μm, n=10 biological replicates. Asterisk=vessel lumen.* B Representative brightfield images of Alizarin Red S positive plaque calcification (top left) and corresponding Wnt5a-positive area (bottom left). *Scale bar 500 μm. Asterisk=vessel lumen. Top right*: Quantification of Wnt5a immunohistochemistry as relative positive area per specimen compared to isotype control. *Mean±SD, Shapiro-Wilk normality test followed by unpaired student’s t-test, n=10 biological replicates. Bottom right*: Correlation of relative calcified lesion area with relative Wnt5a-positive area by linear regression. *Dotted line: 95% confidence interval, Pearson analysis; n=10 biological replicates, n=2 specimen each.* C Representative immunofluorescent images for CD68^+^MMP7^+^S100A8^+^ cells at 20x magnification. *Scale bar 50 μm, n=10 biological replicates. Asterisk=vessel lumen.* D Inlay at 63x magnification. *Scale bar 5 μm.* E Immunofluorescence for CD68^+^pCaMKII^+^ cells from the same specimen at 63x magnification. *Scale bar 5 μm.* F Linear regression of relative calcified lesion area with CD68^+^MMP7^+^S100A8^+^ cells per mm^2^ tissue. **G** Linear regression of relative calcified lesion area (*top*) or CD68^+^MMP7^+^S100A8^+^ cells per mm^2^ tissue (*bottom*) with CD68^+^pCaMKII^+^ cells per mm^2^ tissue. *Pearson analysis with associated two-tailed p-value; dotted line: 95% confidence interval, n=10 biological replicates, n=2 specimen each*.

## Discussion

This study contributes to the characterization of the CXCL4-induced mononuclear cell phenotype with respect to its role in vascular inflammatory processes. Herein, we report a precedential effect of CXCL4 on peripheral blood-derived monocytes causing an induction of osteogenic marker expression concomitant with augmented release of calcific extracellular vesicles and EV-specific enrichment of matrix metalloproteinase-7. Through the aforementioned implications and their dependency on Wnt5a-CaMKII-mediated signaling first identified herein, this distinct subtype of monocytes/macrophages is postulated to propagate extracellular matrix calcification and thus contribute to unfavorable plaque remodeling driving vascular inflammation and atherosclerosis progression.

Emerging evidence on CXCL4-polarized inflammatory effector cells has purported a contributory role in plaque genesis and growth: the CXCL4-induced transcriptome characterized by upregulation of MMP7 and S100A8 first described by Gleissner et al. coincided with a marked suppression of efferocytotic capacity in human PBMCs^5^. At organismal level, knockout of CXCL4 in LDL receptor-deficient mice resulted in a significant reduction of plaque burden compared to single-knockout littermates concurrent with decreased macrophage infiltration, likely owing to attenuated monocyte chemotaxis^29^. In coronary artery plaques from explanted human hearts, the occurrence of MMP7- and S100A8-expressing CD68-positive cells was associated with histological features of plaque instability^30^. Complementary single-cell RNA sequencing data from human coronary artery plaques presented herein confirmed the presence of a CXCL4-susceptible plaque macrophage population bearing a distinct vesicle-directed transcriptome functionally related to macrophage calcium binding capacity. Despite these comprehensive data assuming CXCL4-polarized mononuclear cells as protagonists in progressive plaque inflammation through a putative EV-based effect, the causative mechanisms and effector functions facilitating the CXCL4-dependent monocyte/macrophage phenotype remain nebulous.

S100A8, alternatively termed calgranulin A, is known to form a heterodimer with S100A9 to exert a dual role as calcium-binding protein and potent modulator of inflammation^31^. MMP7, or matrilysin, has previously been implicated in detrimental ECM remodeling through proteoglycan degradation, thus favoring atherosclerotic plaque destabilization^32^. The results presented herein support a novel mechanism of CXCL4-induced EV release and specific enrichment of PBMC-derived EV with S100A8, MMP7 and ALP conducive to enhanced overt calcification *in vitro*, thus potentially promoting adverse ECM transformation under inflammatory conditions. In accordance with *in vitro* data, the present study describes an unprecedented correlation between CD68^+^MMP7^+^S100A8^+^ mononuclear cells commonly associated with the CXCL4-induced phenotype and progressive plaque calcification in human carotid artery explants. With respect to CXCL4-activated signaling in PBMCs, a causal association of CXCL4-dependent gene and protein expression signatures as well as MMP7^+^S100A8^+^ calcific EV release and subsequent overt calcification with noncanonical Wnt5a-Ca^2+^ pathway activation is observed *in vitro*, and a consistent quantitative and spatial correlation of Wnt5a protein expression and CaMKII phosphorylation in CD68^+^ cells with CD68^+^MMP7^+^S100A8^+^ cell abundance and plaque calcification is appreciated in human carotid artery specimen. Based on present evidence, we thus postulate that previously reported effects of CXCL4-polarized mononuclear cells on atherosclerotic plaque progression and destabilization may be footed on an EV-mediated ECM remodeling process involving calcium-phosphate nucleation in the presence of ALP and S100A8 with consecutive degradation of structural ECM components by MMP7 released from calcifying EV. Thus, the evidence provided in this study complements existing data on a platelet-macrophage crosstalk driving pathological ECM remodeling in tissue inflammation^33^.

Additional functional data is needed to corroborate the connection between CXCL4-differentiated monocyte/macrophage-mediated calcific EV release, MMP7 and S100A8 loading and calcification formation. To this end, experimental models may help elucidate the functional contribution of the CXCL4-induced monocyte/macrophage phenotype relative to other immune cell populations to calcific plaque progression. Furthermore, broad spectrum analysis of CXCL4-induced PBMC-derived EV may identify EV-bound factors crucial to cell-cell communication, driving osteogenic/inflammatory phenotype transition of cells in spatial proximity. Lastly, a putative effect of CXCL4-induced PBMC-derived EV on the balance of macro- and microcalcifications, a paramount criterion for plaque stability as a major determinant of rupture risk^34^, needs to be investigated in subsequent studies. Further exploration of the contribution of the CXCL4-polarized monocyte/macrophage phenotype and its associated Wnt5a-regulated, EV-mediated response in vascular inflammation and calcification may pose a promising target of novel, target-specific therapies for plaque destabilization in atherosclerosis, an inflammation-driven disease currently devoid of causative therapies.

## Acknowledgements

The authors thank Prof. Dr. Christian Buchholz, PhD and Dr. Frederic Thalheimer, PhD at Paul Ehrlich Institute in Langen, Germany, for their excellent assistance with Nanoparticle Tracking Analysis. Furthermore, the authors acknowledge PD Dr. med. Andreas Peters of the Vascular Biomaterial Bank Heidelberg (VBBH) for his assistance with human carotid punch specimens.

## Non-standard Abbreviations and Acronyms

PBMC: Peripheral blood-derived mononuclear cell
CXCL4: Chemokine (C-X-C motif) ligand 4
EV: Extracellular vesicle
MMP7: Matrix metalloproteinase 7
S100A8: S100 calcium binding protein A8/calgranulin A
NTA: Nanoparticle tracking analysis
ALP: Alkaline phosphatase
β-GP: β-glycerophosphate
OM: Osteogenic media
M-CSF: Monocyte/macrophage colony stimulating factor
vSMC: Vascular smooth muscle cell
(p)CaMKII: (phospho-)Calcium-calmodulin kinase II

## Sources of Funding

This study was supported by personal funds from the German Centre for Cardiovascular Research (DZHK) awarded to J.B.K.

## Disclosures

None.

## Supplemental Materials

Supplemental Methods

Major Resources Table

Supplemental Figures S1-6

Supplemental Table 1

References 10-16

## Highlights

- CXCL4 induces a pro-inflammatory, osteogenic phenotype in PBMCs characterized by gene and protein expression as well as specific EV enrichment of MMP7 and S100A8
- PBMCs polarized by CXCL4 exhibit augmented secretion of EV bearing the potential to calcify extracellular matrix and illicit a pro-inflammatory response in proximate vascular smooth muscle cells
- The CXCL4-induced PBMC phenotype and calcific EV release show dependency of paracrine Wnt5a-CaMKII signaling
- EV released by CXCL4-induced monocytes/macrophages propagate calcification of the extracellular matrix *in vitro*, and CXCL4-induced mononuclear cell abundance associates with plaque calcification in human carotid artery specimen

## Notes

### Competing Interest Statement

HAK: personal fees from AstraZeneca, Bayer Vital, Daiichi Sankyo, Boehringer Ingelheim, Roche Diagnostics.

