## Supplementary material for "CXCL4-induced PBMCs Modulate Vascular Extracellular Matrix through Wnt5a-CaMKII-dependent Release of Calcific Extracellular Vesicles and Matrix Metalloproteinase-7": Data Supplement

#### Supplemental Methods

*PBMC isolation and cell culture*

Fresh blood samples were obtained from healthy donors of both sexes following approval by the Institutional Review Board at the Faculty of Medicine at Heidelberg University (IRB #S123/2022; German Registry for Clinical Studies #DRKS00029695), and prior written consent was obtained from all donors. All methods were performed in accordance with the Declaration of Helsinki and its later amendments as well as local and national guidelines and regulations. Following asservation of up to 200 ml of whole blood from the antecubital vein, isolation of peripheral blood mononuclear cells (PBMCs) was essentially done as described previously[10, 11] with slight modifications. Briefly, blood was drawn into ethylenediamine tetraacetate (EDTA) containing tubes to a final concentration of 10 mM, and buffy coats were separated immediately by density gradient centrifugation using FicoLite-H 1.077 g/ml (Linaris, Dossenheim, Germany). After washing, monocytes were isolated using a magnetic bead-based CD14 negative selection kit according to manufacturer's instructions (STEMCELL Technologies, Vancouver, Canada). Isolated PBMCs were seeded out at $10^5$ cells per $cm^2$ and differentiated in serum-free Macrophage-SFM (Life Technologies, Carlsbad, USA) supplemented with 1% Nutridoma (Roche, Basel, Switzerland), 100 IE/ml penicillin, 100 $\mu$g/ml streptomycin (Life Technologies) and either 100 ng/ml recombinant human M-CSF as control or 1 $\mu$M recombinant human CXCL4 (Peprotech, Hamburg, Germany). Media was changed every 48 hours.
For experiments involving osteogenic induction, differentiation media was switched to osteogenic differentiation media containing 10 mM $\beta$-glycerophosphate (Merck, Darmstadt, Germany) and 2,6 mM $Ca^{2+}$ (Roth, Karlsruhe, Germany) for 24 hours before harvest. For Wnt5a stimulation, differentiation media was switched to osteogenic differentiation media, as described above, additionally supplemented with either 250 ng/ml recombinant human Wnt5a (R&D Systems, Minneapolis, MN, USA) or 250 ng/ml rhWnt5a + 100 $\mu$M Box5 (Merck, Darmstadt, Germany), a Wnt5a-derived hexapeptide antagonist of Wnt5a-mediated auto-/paracrine signaling[12], for 24 hours before harvest.

*Extracellular vesicle (EV) isolation*

Extracellular vesicles were essentially isolated from cell culture media as described previously[13-15]. Briefly, PBMCs were cultured in differentiation media as detailed above for 6 days. On day 6, media was changed to either differentiation media, osteogenic differentiation media or osteogenic differentiation media supplemented with rhWnt5a or rhWnt5a+Box5. After 24 hours in culture, media supernatant was harvested and deprived of cell debris and larger apoptotic bodies by incremental centrifugation steps. Lastly, media EV fraction was pelleted by ultracentrifugation at $10^5$ x *g* at 4°C overnight on an Optima XPN-80 centrifuge with an SW40 Ti rotor (Beckman Coulter, Brea, CA, USA) and resuspended accordingly depending on downstream analysis. Pellets were either analyzed immediately or stored at -80°C until further analysis. An equal volume of cell-free culture media was treated in parallel to serve as media control.

*Nanoparticle Tracking Analysis (NTA)*

For EV quantification and size distribution analysis, the EV pellet was resuspended in 100 μl phosphate buffered saline (PBS) to create an EV stock solution. For each sample, working dilutions were titrated to achieve a final rate of 10-50 particles per frame on nanoparticle tracking. Diluted EV samples were visualized on a Nanosight NS300 (Malvern Panalytical, Malvern, UK) using a syringe pump at a constant flow rate of 40 μl/min. For each sample, 4 separate runs of 30s each were recorded, and for analysis of Brownian motion of particles in solution Nanosight NTA software v3.3 at a camera level of 16 and detection threshold of 9 was used. Size and concentration data from each recorded run were averaged to yield values representative for each sample.

*In vitro EV alkaline phosphatase (ALP) activity assay*

Isolated EV pellets were lysed in ALP assay buffer containing 0.2% (v/v) Triton X-100. After incubation for 10 min at 4°C under gentle agitation, ALP activity was quantified using the SensoLyte pNPP Colorimetric Assay Kit (Anaspec, Fremont, CA, USA) according to manufacturer's instructions. Absorbance was read at 405 nm in duplicates, and ALP activity was calculated from a standard curve. Activity values were normalized to EV protein content determined by bicinchoninic acid (BCA) assay (Thermo Fisher, Waltham, MA, USA), and averaged activity was further normalized to M-CSF-differentiated control.

*Alizarin Red S staining*

PBMCs cultured in 24-well plates were fixed in 4% (v/v) paraformaldehyde for 20 min at room temperature. Following washing with PBS, cells were incubated with 250 μl per well of Alizarin Red S (ARS) solution (Morphisto, Offenbach, Germany) for 5 mins at room temperature. After aspiration and washing to remove excess dye, brightfield images of ARS staining were taken with an EVOS XL Core microscope (Life Technologies). For quantification of staining intensity, complexed dye was dissolved in 100 mM cetylpyridinium chloride (Merck) for one hour under gentle agitation, and absorbance of resulting solutions was read at 550 nm in duplicates.
Frozen tissue sections were thawed at 4°C, fixed with 4% paraformaldehyde in PBS for 20 mins at 4°C and rehydrated in PBS at room temperature, followed by ARS solution for 5 mins. Dyed sections were rinsed briefly in acetate buffer (pH 4.0) and dehydrated in 96% ethanol and isopropanol. Images were obtained on Olympus BX51 and Leica MZFLIII brightfield microscopes. For staining quantification, positive area was measured using ImageJ and divided by total tissue area. For each biological replicate, two tissue sections were stained.

*EV-vSMC coculture*

EV isolates from differentially stimulated PBMCs obtained from n=3 biological replicates were pooled and purified by centrifugal filtration using Amicon Ultra Centrifugal Filter Units (Millipore, Burlington, MA, USA; nominal molecular weight cutoff of 10 kDa). Protein content of EV samples was determined by BCA assay (Thermo Fisher), and equal amounts of EV were used for coculture experiments.
Human vascular smooth muscle cells (vSMC) were purchased from PromoCell. VSMC between passages 5 and 9 were seeded out on to 6-well plates at $3*10^4$ cells/cm$^2$ and were subsequently allowed to settle for 48 hours in DMEM supplemented with 10% (v/v) fetal calf

serum, 100 IE/ml penicillin and 100 μg/ml streptomycin (all purchased from Life Technologies). Hereafter, vSMC were starved in DMEM with 1% (v/v) exosome-depleted fetal calf serum and antibiotics supplemented with either 10 mM β-glycerophosphate/2,6 mM $Ca^{2+}$ (osteogenic media), vehicle or PBMC-derived EV. Exosome depletion of fetal calf serum was achieved through ultracentrifugation at $10^5$ x *g* at 4°C overnight. After 24 hours in conditioned media, vSMC were harvested for subsequent analysis.

*EV fluorescent labeling*

To investigate selective uptake of PBMC-derived EVs by vSMC, EVs were labeled using the green fluorescent lipophilic dye PKH67 (Sigma Aldrich, St. Louis, MO, USA) essentially as described previously[16], with slight modifications. Briefly, PBMC-derived EV isolates were diluted to 1 ml, and 6 μl of PKH67 dye concentrate was added. After gentle mixing and incubation for 5 mins, solutions were quenched with 10% bovine serum albumin (Serva, Heidelberg, Germany) in PBS and further diluted in DMEM (Life Technologies). To separate labeled EV from excess dye, a 0.971 M sucrose phase was carefully added to the bottom of each tube, and labeled EV were pelleted at $1.9*10^5$ *x g* for 2 hours at 4°C. The resulting EV pellet was washed using Amicon Ultra Centrifugal Filter Units (Millipore; nominal molecular weight cutoff of 10 kDa) according to manufacturer's instructions. An equal volume of PBS was treated in parallel to serve as vehicle control.
Resulting EV concentrates or vehicle control were resuspended in vSMC starvation medium (consisting of DMEM with 1% exosome-depleted fetal calf serum and antibiotics) and added to vSMC that were seeded out on to 8-well chamber slides ($5*10^4$/well) 48 hours prior. After 24 hours of culture, vSMC were washed in PBS and fixed with 4% (v/v) paraformaldehyde for 20 mins at room temperature. Following additional washing steps, cell nuclei were counterstained with 4',6-Diamidino-2-phenylindol (DAPI; Life Technologies) diluted $1:10^4$ for one minute in the dark. PKH67 fluorescence at 488 nm was detected using an Axio Observer Z1 fluorescence microscope (Zeiss, Oberkochen, Germany). For fluorescence quantification, representative images from n=2 technical replicates per biological replicate were taken. For each separate image, fluorescent area was quantified using ImageJ and divided by the number of nuclei to yield average fluorescent intensity per cell. Measurements were averaged and compared to vehicle control.

*Quantitative real-time Polymerase Chain Reaction (qRT-PCR)*

Cultured cells were harvested in buffer RLT (Qiagen, Hilden, Germany) supplemented with 500 μM β-Mercaptoethanol (Life Technologies) using a rubber policeman. Cell lysates were homogenized with QiaShredders, and total RNA was isolated with the RNeasy Mini Kit (Qiagen) according to manufacturer's instructions. RNA content was quantified on a DS11 spectrophotometer (DeNovix, Wilmington, DE, USA). Reverse transcription was performed with the iScript cDNA Synthesis Kit on a C1000 Thermal Cycler (Bio-Rad, Hercules, CA, USA). Messenger RNA expression levels were determined by SYBR Green-based quantitative real-time PCR (Bio-Rad) on a Quant Studio 5 system (Life Technologies). Human primers were purchased from Eurofins Genomics, a list of primers used for analysis is included in the major resources table. Expression levels were normalized to GAPDH and M-CSF-differentiated control using the ΔΔCT method and presented as fold increase relative to control.

*Western Blot*

For isolation of protein from cultured cells, flow-through from RNA isolation was collected, and protein was precipitated in acetone at -20°C and resuspended in radio-immunoprecipitation assay (RIPA) buffer (Abcam, Cambridge, UK) supplemented with protease and phosphatase inhibitors (Cell Signaling, Danvers, MA, USA) as per manufacturer's instructions. For EV protein isolation, EV pellets were lysed in appropriate

volume of RIPA buffer with protease/phosphatase inhibitors. Protein lysates were homogenized by brief sonication (Diagenode, Seraing, Belgium), and protein content was quantified by BCA assay. Equal amounts of protein were separated by 4-15% sodium dodecylsulfate polyacrylamide gel electrophoresis (SDS-PAGE) (Bio-Rad) and subsequently transferred to polyvinylidene difluoride (PVDF) membranes (Millipore) using the Bio-Rad Mini Trans-Blot system. After unspecific protein blocking with 3% (w/v) bovine serum albumin (BSA) diluted in tris-buffered saline with 0.5% (v/v) Tween-20 (Bio-Rad) for one hour, membranes were incubated with primary antibody diluted in blocking buffer overnight at 4°C under gentle agitation. Following incubation with species-matched horse radish peroxidase (HRP)-conjugated secondary antibody, protein expression was detected using the SuperSignal West Femto Maximum Sensitivity Substrate (Thermo Fisher) on a Fusion FX imaging system (Vilber Lourmat, Marne-la-Vallée, France). Relative band intensity was calculated and normalized to housekeeping protein as well as M-CSF-differentiated control using NIH ImageJ version 1.53. To quantify expression of phosphorylated proteins, total protein and phosphorylated protein were first separately normalized to housekeeping protein, after which a ratio of phosphorylated/total protein was calculated. Ratios of all experimental groups were normalized to control.
A list of all antibodies used for analysis is found in the major resources table.

*Human carotid artery tissue*

Carotid artery punch specimens (n=10) from patients undergoing carotid thromboendarterectomy surgery at University Hospital Heidelberg were obtained from the Vascular Biomaterial Bank Heidelberg (VBBH), a sub-repository of the BioMaterial Bank Heidelberg (BMBH). All patients had given informed written consent prior to surgery, and all sample collections for the VBBH had received prior approval by the Institutional Review Board at the Medical Faculty of Heidelberg University (IRB #S-149/2010, S-301/2013 and amendment of 2016). Separate approval was obtained for the use of tissue specimens in this study (IRB #584/2022). Besides age (73.3 ±8.8 years) and sex (90% male), no additional demographic or clinical information was made available to the authors prior to analysis. Frozen carotid tissue specimens were cut into 10 μm thick consecutive sections (two sections per slide) and stored at -80°C until further analysis.

*Immunohistochemistry*

Carotid punch tissue sections were thawed at 4°C, fixed with 4% paraformaldehyde in PBS for 20 mins at 4°C and rehydrated in PBS at room temperature. Following quenching of endogenous peroxidase activity with 3% (v/v) hydrogen peroxide in methanol, sections were blocked in 5% BSA in PBS for 1 hour. Subsequently, primary antibody or species- and concentration-matched isotype control (Becton Dickinson, Franklin Lakes, NJ, USA) diluted in 2.5% (w/v) BSA/PBS with 0.1% Triton X-100 were incubated overnight at 4°C, followed by appropriate secondary antibody for one hour at room temperature. Staining was visualized using the HRP/DAB detection kit (Abcam) according to manufacturer's instructions. Weigert's iron hematoxylin was used as nuclear counterstain (Sigma Aldrich). Representative and overview brightfield images were obtained on an Olympus BX51 and Leica MZFLIII stereomicroscope, respectively.

*Immunofluorescence – cells*

PBMCs were seeded out on to 4-well chamber slides ($5*10^4/cm^2$) and differentiated accordingly. Cell fixation was achieved with 4% paraformaldehyde for 30 mins at room temperature, followed by permeabilization with 0.25% (v/v) Triton X-100 for one hour. Nonspecific protein binding was blocked with 3% (w/v) BSA in PBS for one hour. Primary antibody diluted in 0.1% (w/v) BSA/PBS was incubated overnight at 4°C, followed by

appropriate fluorophore-conjugated secondary antibody for one hour at room temperature. Nuclei were counterstained with DAPI (Life Technologies) 1:$10^4$ for one minute. Secondary antibody-only staining served as negative control. Images were obtained on a Zeiss Axio Observer Z1 fluorescence microscope.
For staining quantification, representative overview images from n=3 technical replicates per sample were taken. For each image, percentage of positively stained area was determined using ImageJ and divided by the number of DAPI-stained nuclei. Relative positive area was averaged for each sample and subsequently normalized to M-CSF stimulated control.

*Immunofluorescence – carotid artery tissue*

Frozen tissue sections were thawed, fixed and hydrated as described above. Unspecific protein blocking was achieved with 5% (w/v) BSA/PBS for one hour at 4°C. Sections were subsequently incubated with primary antibody or concentration-matched rabbit (Becton Dickinson) or mouse (Abcam) isotype control diluted in 2.5% BSA/PBS +0.1% Triton X-100 overnight at 4°C. Following washing steps, biotinylated anti-polyvalent secondary antibody was added for one hour at room temperature, followed by fluorophore-conjugated streptavidin diluted in 2.5% BSA/PBS for 30 mins. For double/triple immunofluorescence, all steps starting with primary antibody were repeated once/twice, each time preceded by an avidin/biotin blocking step (GeneTex, Irvine, CA, USA) to avoid fluorescent labeling of unbound biotin from a previous step. Lastly, sections were counterstained with DAPI (Life Technologies) diluted 1:$10^4$ in PBS for one minute and mounted in FluorSave reagent (Millipore). Two sections per biological replicate were stained. Images were obtained on a Leica SP8 confocal microscope. For each tissue section, positive cells were counted using the Leica Application Suite X software version 3.5.7, and the number of positive cells was divided by total tissue area.
The following sequences of antibodies and fluorophores were used (detailed information on each antibody can be found in the major resources table):
*Double immunofluorescence:*
Mouse-anti-pCaMKII (Abcam) – goat-anti-polyvalent biotinylated secondary antibody (Abcam) – streptavidin-Texas Red (Vector Laboratories, Newark, CA, USA) – rabbit-anti-CD68 (Cell Signaling) – goat-anti-polyvalent biotinylated secondary antibody (Abcam) – streptavidin-Alexa Fluor 488 (Thermo Fisher)
*Triple immunofluorescence:*
Mouse-anti-S100A8 (BMA Biomedicals) – goat-anti-polyvalent biotinylated secondary antibody (Abcam) – streptavidin-Alexa Fluor 647 (Thermo Fisher) – rabbit-anti-CD68 (Cell Signaling) – goat-anti-polyvalent biotinylated secondary antibody (Abcam) – streptavidin-Texas Red (Vector Laboratories) – rabbit-anti-MMP7 (Abcam) – goat-anti-polyvalent biotinylated secondary antibody (Abcam) – streptavidin-Alexa Fluor 488 (Thermo Fisher)

*Picrosirius Red staining*

Tissue sections were thawed, fixed and hydrated as described above. Sections were stained in saturated aqueous picric acid solution (Morphisto) for one hour, rinsed briefly with 30% (v/v) acetic acid and dehydrated with ethanol and isopropanol before xylene clearing and mounting. Weigert's iron hematoxylin was used as nuclear counterstain. Images under polarized light were obtained on a Zeiss Axio Imager M2.

*Analysis of single-cell RNA sequencing data from human coronary artery plaques*

A human single-cell RNA sequencing repository from plaque tissue extracted from the right coronary artery of n=4 heart transplant recipients was accessed through the Gene Expression Omnibus database (GSE131778). Data analysis was conducted using the R package Seurat version 4.3.0. For quality control purposes, cells expressing <500 or >3500 genes or >7.5%

mitochondrial genes were excluded from analysis. Likewise, genes expressed in fewer than five cells were excluded. Raw gene expression values were log-normalized with a scale factor of $10^4$. For subsequent analyses, the most highly variable genes were selected using the variance-stabilizing transformation method. Principal component analysis was conducted for linear dimension reduction, and clusters were identified by a shared nearest neighbor (SNN) modularity optimization-based algorithm. The t-distributed stochastic neighbor embedding (t-SNE) tool was utilized to visualize resulting clusters in a two-dimensional roster. Following comparative gene expression signature-based identification of cell clusters, a cell subset comprising only monocyte-derived macrophages (MDM) and resident macrophages was extracted, and cluster identity was reassigned dichotomously based on *CCR1* expression. Gene Ontology (GO) database annotations for cell component and molecular function were identified by enrichment analysis of the top 100 differentially expressed genes in *CCR1*$^+$ plaque mononuclear cells using the *Enrichr* tool.

*Statistical Analysis*

Statistical analyses were performed using GraphPad Prism version 9.0.0. Shapiro-Wilk test was applied to test for normality. For normally distributed parameters, student's t-test was used for comparison between two groups. For not normally distributed parameters, a Mann-Whitney U test was applied. For three or more experimental groups, analysis of variance (ANOVA) or mixed effects analysis with Bonferroni post-hoc test for multiple comparisons was applied for normally distributed parameters, where appropriate. For non-normally distributed parameters, Kruskal-Wallis test with Dunn's multiple comparison test was applied. Error bars represent standard deviation, unless otherwise specified. A p-value of 0.05 was defined to denote statistical significance.
For single-cell RNA sequencing analyses, differentially expressed genes were identified using Wilcoxon's rank sum test implemented through the R Seurat package. Gene Ontology (GO) annotation-based enrichment analysis was performed using Fisher's exact test with Benjamini-Hochberg adjustment for multiple testing, and ranking was done based on resulting p-values.
For correlation analyses in Fig. 7, linear regression was performed, and Pearson's $R^2$ was calculated. To test for significance of correlation, a t-test was performed assuming $H_0$: correlation coefficient $\rho = 0$ and $\alpha = 0.05$.

### Major Resources Table

qRT-PCR Primers:

| target | forward (5’ – 3’) | reverse (5’ – 3’) |
|---|---|---|
| GAPDH | *GGCTCATGACCACAGTCCAT* | *GCCTGCTTCACCACCTTCT* |
| MMP7 | *GAGTGAGCTACAGTGGGAACA* | *CTATGACGCGGGAGTTTAACAT* |
| S100A8 | *ATGCCGTCTACAGGGATGAC* | *ACTGAGGACACTCGGTCTCTA* |
| ALP | *GACCCTTGACCCCCACAAT* | *GCTCGTACTGCATGTCCCCT* |
| OPN/SPP1 | *GCCGAGGTGATAGTGTGGTT* | *TGAGGTGATGTCCTCGTCTG* |
| RUNX2 | *CCAACCCACGAATGCACTATC* | *TAGTGAGTGGTGGCCGACATA* |
| CTNNB1 | *GCTTTCAGTTGAGCTGACCA* | *CAAGTCCAAGATCAGCAGTCTC* |
| WNT1 | *CGCTGGAACTGTCCCACT* | *AACGCCGTTTCTCGACAG* |
| WNT2 | *TTTGGCAGGGTCCTACTCC* | *CCTGGTGATGGCAAATACAA* |
| WNT2B | *AACTTACATAATAACCGCTGTGGTC* | *ACTCACGCCATGGCACTT* |
| WNT3 | *CTCGCTGGCTACCCAATTT* | *GAGCCCAGAGATGTGTACTGC* |
| WNT3A | *AACTGCACCACCGTCCAC* | *AAGGCCGACTCCCTGGTA* |
| WNT4 | *GCAGAGCCCTCATGAACCT* | *CACCCGCATGTGTGTCAG* |
| WNT5A | *ATTGTACTGCAGGTGTACCTTAAAAC* | *CCCCCTTATAAATGCAACTGTTC* |
| WNT5B | *GGAGCGAGAGAAGAACTTTGC* | *CGTCGTCCATCTTATACACAGC* |
| WNT6 | *AGAGTGCCAGTTCCAGTTCC* | *GAACACGAAGGCCGTCTC* |
| WNT7A | *CTTCGGGAAGGAGCTCAAA* | *GCAATGATGGCGTAGGTGA* |
| WNT7B | *CGCCTCATGAACCTGCATA* | *GCTGCATCCGGTCCTCTA* |
| WNT8A | *GTGATGGGTCAAACAATGGA* | *ATCCTTTCCCCAAATTCCAC* |
| WNT8B | *TGTGATGACTCCCGCAAC* | *CGAAGCCCACATTGTCACT* |
| WNT9A | *TCCAGTTCCGCTTTGAGC* | *AGCCGAGGAGATGGCATAG* |
| WNT9B | *GCCTCCCTCGATACTCAACA* | *ACAAGGTTGGGGATGCTTG* |
| WNT10A | *ATCCACGCGAGAATGAGG* | *CCGCATGTTCTCCATCACT* |
| WNT10B | *ATGCGAATCCACAACAACAG* | *TCCAGCATGTCTTGAACTGG* |
| WNT11 | *AGCTCGCCCCCAACTATT* | *ATACACGAAGGCCGACTCC* |
| WNT16 | *CAATGAACCTACATAACAATGAAGC* | *CAGCGGCAGTCTACTGACAT* |
| CAMKIIA | *ACCACTACCTGATCTTCGACC* | *CCGCCTCACTGTAATACTCCC* |
| IL6 | *ACTCACCTCTTCAGAACGAATTG* | *CCATCTTTGGAAGGTTCAGGTTG* |

Primary Antibodies:

| target | species | company | catalog # | application |
|---|---|---|---|---|
| gapdh | rabbit | Abcam | ab181602 | WB |
| s100a8 | mouse | Santa Cruz | sc-48352 | WB |
| s100a8 | mouse | BMA Biomedicals | T-1030 | IF |
| annexin V | rabbit | Cell Signaling | 8555 | WB |
| mmp7 | rabbit | Abcam | ab207299 | WB, IF |
| cd9 | rabbit | Cell Signaling | 13403 | WB |
| active β-catenin | rabbit | Abcam | ab246504 | WB |
| nonactive (phospho)-β-catenin | rabbit | Abcam | ab223075 | WB |
| pCaMKII (Thr286) | mouse | Abcam | ab171095 | WB, IF |
| CaMKII (total) | rabbit | Abcam | ab92332 | WB |
| pp38 (Thr180/Tyr182) | rabbit | Cell Signaling | 9211 | WB |
| p38 (total) | rabbit | Cell Signaling | 9212 | WB |
| pJNK 1/2/3 | rabbit | Abcam | ab76572 | WB |
| JNK 1/2/3 (total) | rabbit | Abcam | ab179461 | WB |
| pp44/42 MAPK (Erk 1/2) | rabbit | Cell Signaling | 4376 | WB |
| Erk 1/2 (total) | rabbit | Cell Signaling | 9102 | WB |
| wnt5a | rabbit | Abcam | ab227229 | WB |
| wnt5a | rabbit | Abcam | ab235966 | IHC |
| cd68 | rabbit | Cell Signaling | 76437 | IF |

Secondary Antibodies:

| target (conjugate) | species | company | catalog # | application |
|---|---|---|---|---|
| anti-rabbit (HRP) | goat | Cell Signaling | 7074 | WB |
| anti-rabbit (HRP) | goat | Abcam | ab6721 | IHC |
| anti-mouse (HRP) | rabbit | Abcam | ab6728 | WB |
| anti-rabbit (Alexa Fluor 488) | donkey | Abcam | ab150073 | IF |
| anti-rabbit (Alexa Fluor 594) | goat | Abcam | ab150080 | IF |
| anti-rabbit/mouse (biotinylated) | goat | Abcam | ab64264 | IF |

### Supplemental Table 1

| Wnt | Pathway | IFNγ+LPS/CXCL4 versus M-CSF polarization |
|---|---|---|
| 1 | *Canonical* | Variable expression, partly below detection limits |
| 2 | *Canonical* | Variable expression, partly below detection limits |
| 2B | *n/d* | Mildly overexpressed after IFNγ+LPS stimulation |
| 3 | *Canonical* | Variable expression close to detection limits |
| 3A | *Canonical* | No difference in M-CSF/IFNγ+LPS/CXCL4-polarized PBMC |
| 4 | *Wnt-$Ca^{2+}$* | Variable expression close to detection limits |
| **5A** | ***Non-canonical/Wnt-$Ca^{2+}$*** | **Overexpressed in IFNγ+LPS/CXCL4-polarized PBMC** |
| 5B | *Wnt-$Ca^{2+}$* | No difference in M-CSF/IFNγ+LPS/CXCL4-polarized PBMC |
| 6 | *Wnt-$Ca^{2+}$* | No difference in M-CSF/IFNγ+LPS/CXCL4-polarized PBMC |
| 7A | *Wnt-$Ca^{2+}$* | Variable expression close to detection limits |
| 7B | *Wnt-$Ca^{2+}$* | Variable expression, partly below detection limits |
| 8A | *Canonical* | Variable expression, partly below detection limits |
| 8B | *Canonical* | Variable expression, partly below detection limits |
| 9A | *n/d* | Variable expression, no difference in M-CSF/IFNγ+LPS/CXCL4-polarized PBMC |
| 9B | *n/d* | Variable expression, no difference in M-CSF/IFNγ+LPS/CXCL4-polarized PBMC |
| 10A | *Canonical* | Variable expression close to detection limits |
| **10B** | ***Canonical*** | **Overexpressed in IFNγ+LPS/CXCL4-polarized PBMC** |
| 11 | *Non-canonical/Wnt-$Ca^{2+}$* | Mildly overexpressed in CXCL4-polarized PBMC |
| 16 | *n/d* | Variable expression, partly below detection limits |

**Supplemental Table 1**: Summary of gene expression analysis of common Wnt proteins in proinflammatory IFNγ+LPS-stimulated PBMCs and CXCL4-induced PBMCs, compared to M-CSF-stimulated control.

### Supplemental Figure S1

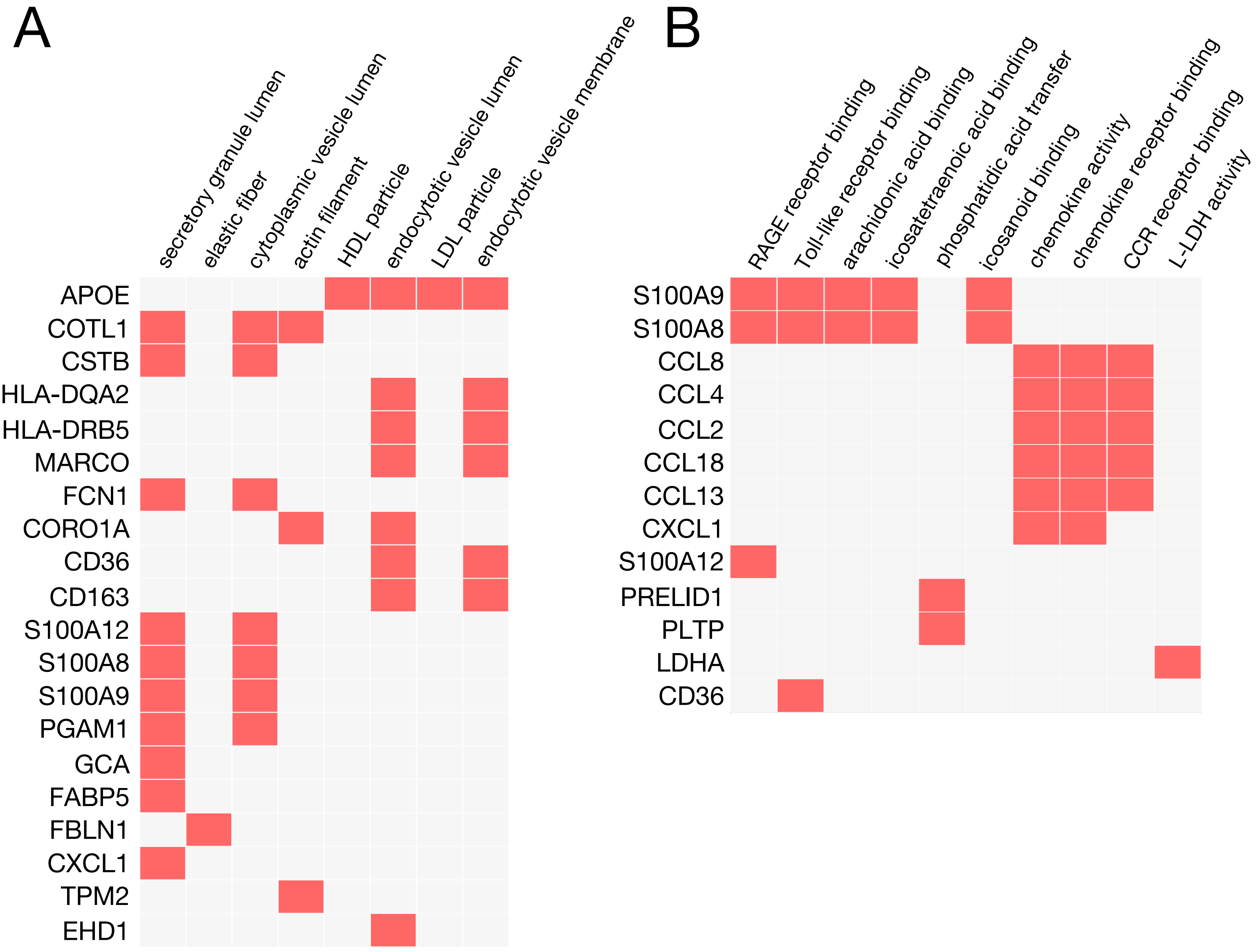


**Figure S1:**. Functional analysis of Gene Ontology (GO) annotations cellular component (**A**) and molecular function (**B**) of top-100 differentially expressed genes in CCR1$^{+}$ mononuclear cells from human coronary plaque tissue using the Enrichr tool. *Data extracted from single cell RNA sequencing repository GEO131778, n=4 biological replicates. Fisher's exact test with Benjamini-Hochberg adjustment.*

### Supplemental Figure S2

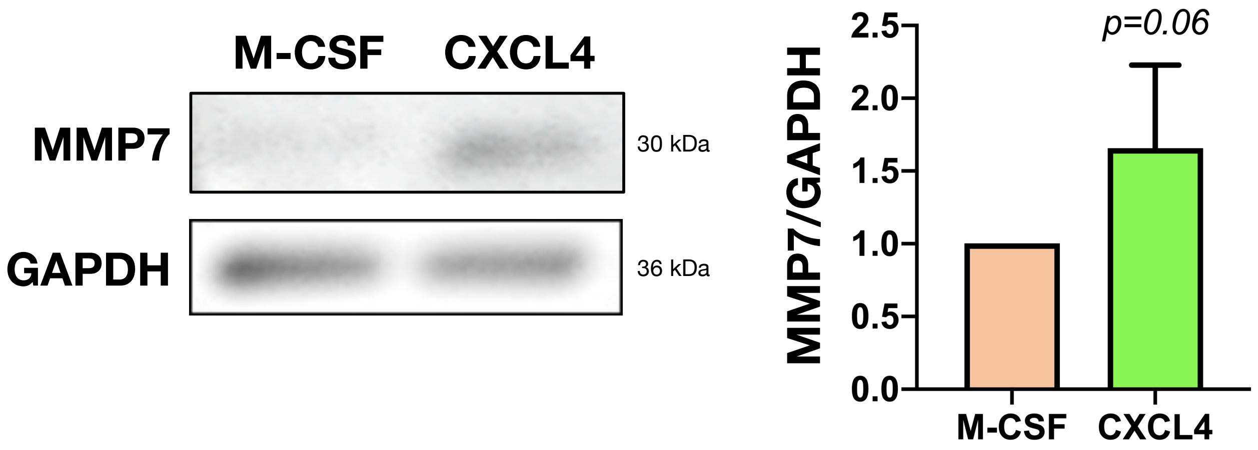


**Figure S2:** Representative immunoblots for MMP7 and GAPDH in whole cell lysates of M-CSF- and CXCL4-differentiated PBMCs (*left*) and Western Blot quantification normalized to M-CSF-stimulated control (*right*). *Mean±SD, Shapiro-Wilk normality test followed by unpaired student's t-test, n=4 biological replicates.*

### Supplemental Figure S3

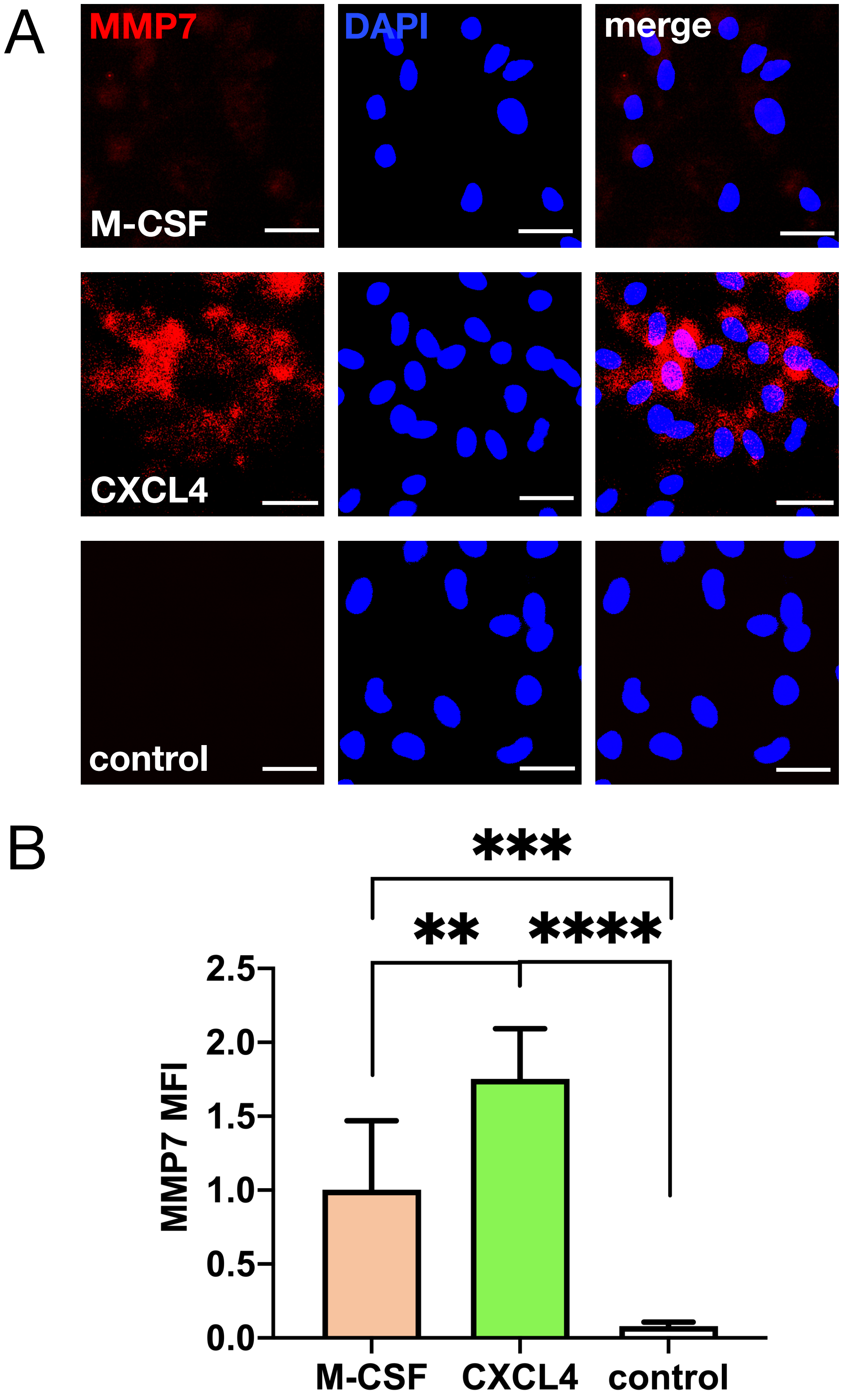


**Figure S3: A** Representative immunofluorescent images for MMP7 in M-CSF- (*top panel*) and CXCL4-differentiated PBMCs (*center panel*) at 40x magnification. *Scale bar 20 μm, n=3 biological replicates.* **B** Quantification of MMP7 mean fluorescence intensity (MFI) normalized to M-CSF-stimulated PBMCs. *Mean±SD, Shapiro-Wilk normality test followed by one-way ANOVA with Bonferroni post-hoc test; n=3 biological replicates. **p<0.01 ***p<0.001 ****p<0.0001.*

### Supplemental Figure S4

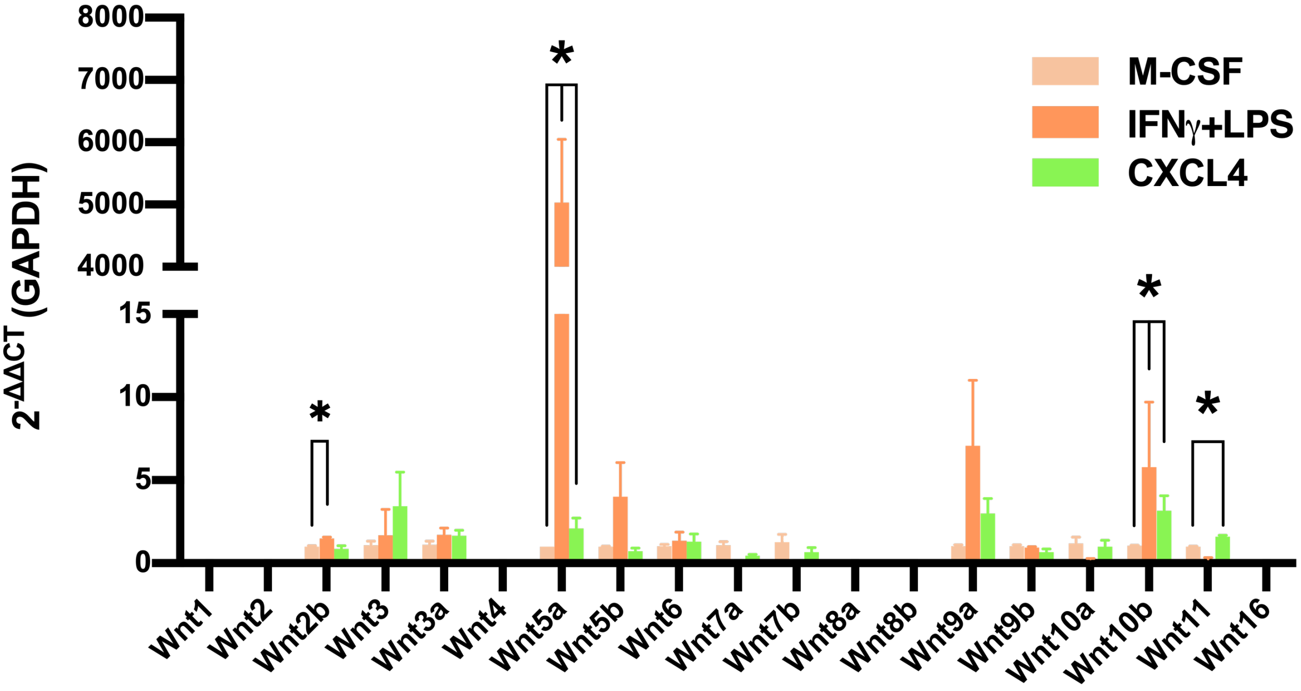


**Figure S4:** Relative mRNA expression of common Wnt proteins in CXCL4- and IFNγ +LPS-stimulated pro-inflammatory PBMCs *in vitro*, normalized to M-CSF-differentiated control. *Mean±SEM, one-way ANOVA; n=3-4 biological replicates. *p<0.05.*

### Supplemental Figure S5

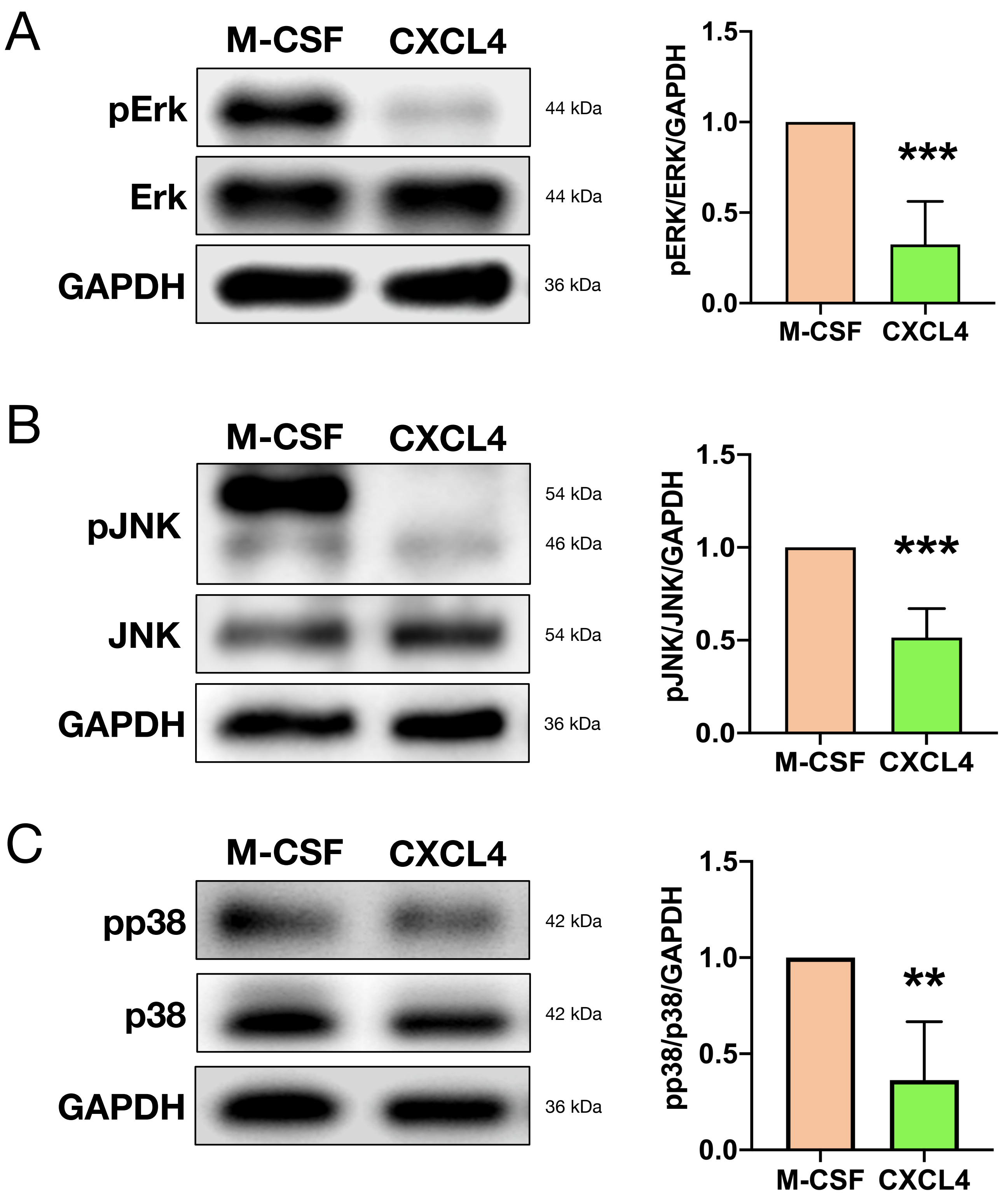


**Figure S5:** Representative immunoblots for **A** phospho-Erk (pErk) **B** phospho-JNK (pJNK) and **C** phospho-p38 (pp38) from whole cell lysates of M-CSF- or CXCL4-differentiated PBMCs, with Western Blot quantification normalized to total protein, GAPDH as loading control and M-CSF-differentiated PBMCs. *Mean±SD, Shapiro-Wilk normality test followed by unpaired student's t-test; n=4 biological replicates. **p<0.01 ***p<0.001.*

### Supplemental Figure S6

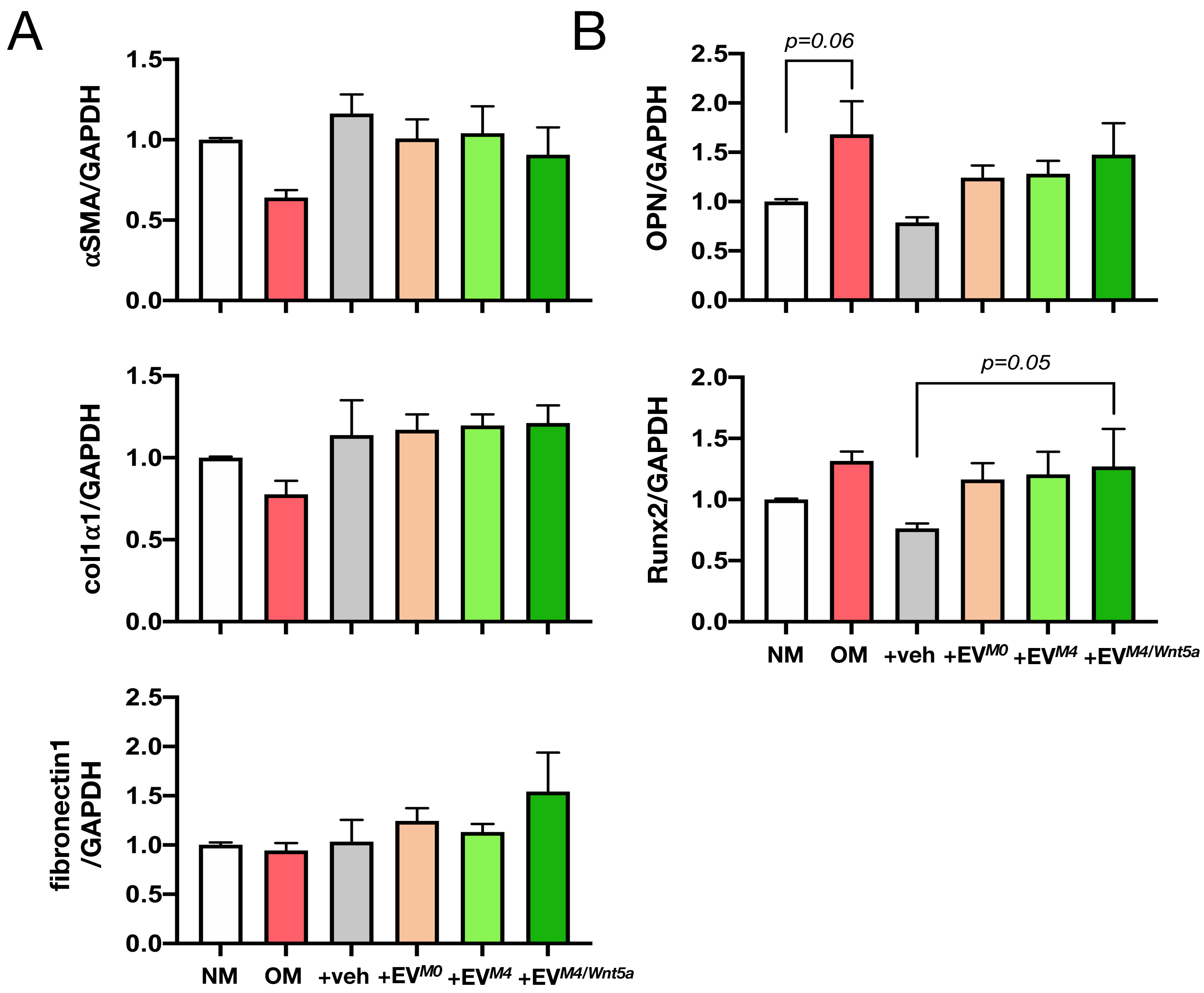


**Figure S6:** Relative mRNA expression of **A** phenotypic markers alpha smooth muscle actin (αSMA), collagen type 1 (col1α1), fibronectin 1, and **B** osteogenic markers osteopontin (OPN) and Runx2 in vascular smooth muscle cells (vSMC) after stimulation with osteogenic media (*OM*), vehicle control (*veh*), EV from M-CSF- ($EV^{M0}$), CXCL4- ($EV^{M4}$) or CXCL4 and Wnt5a-differentiated PBMCs ($EV^{M4/Wnt5a}$), normalized to normal media control (*NM*). *Mean±SEM, Shapiro-Wilk normality test followed by mixed-effects analysis with Bonferroni post-hoc test, n=4 vSMC lines with n=3 PBMC donors, respectively.*
